# Designing a protein with emergent function by combined *in silico, in vitro* and *in vivo* screening

**DOI:** 10.1101/2023.02.16.528840

**Authors:** Shunshi Kohyama, Béla P. Frohn, Leon Babl, Petra Schwille

**Author notes:** These authors contributed equally to this work (Shunshi Kohyama, Béla P. Frohn).

## Abstract

Recently, utilization of machine learning (ML) based methods has led to astonishing progress in protein design and, thus, the design of new biological functionality. However, emergent functions that require higher-order molecular interactions, such as the ability to self-organize, are still extremely challenging to implement. Here, we describe a comprehensive *in silico, in vitro*, and *in vitro* screening pipeline (i^3^-screening) to develop and validate ML-designed artificial homologs of a bacterial protein that confers its role in cell division through the emergent function of spatiotemporal pattern formation. Moreover, we present complete substitution of a wildtype gene by an ML-designed artificial homolog in *Escherichia coli*. These results raise great hopes for the next level of synthetic biology, where ML-designed synthetic proteins will be used to engineer cellular functions.

## Main

The design of novel proteins to perform specific desired functions is one of the ultimate goals of synthetic biology, with the potential to tackle some of the biggest challenges of mankind, including disease, climate change, food, or energy production (*1*–*3*). In the last two years, the introduction of Machine Learning (ML) based generative models, inspired by revolutionary deep generative models like ChatGPT, Stable Diffusion or DallE2 (*4*–*6*), has yielded major breakthroughs in protein design and engineering (*2, 3, 7*–*13*). These methods have led to great advances in the engineering of new proteins with individual functionality, such as catalytic activity (*8*–*11*), small molecule binding (*8, 9*), or spike protein capping (*8*). However, the design of proteins with emergent functions of core relevance to life, such as spatiotemporal self-organization upon energy dissipation, is still in its infancy. Such complex biological functions only arise from finely tuned interactions with the cellular environment, such as other proteins and lipid membranes, and the ability to switch between different conformational states, which is not yet possible to design *de novo* by ML-based design (*14*).

There are two major challenges for this next step of protein engineering. First, there is no comprehensive classification of emergent protein functionalities, quantitatively describing complex protein-protein, -lipid, and -nucleotide interactions within the cellular environment. Thus, there are no good datasets to train generative models on towards *de novo* design of proteins with emergent behavior. Second, there is a lack of screening pipelines, as required for protein design (*2*), to test for emergent functions of *de novo* designed proteins, both computational and experimental. Computational models to predict emergent functions, or the generic biological role of a protein, from sequence or structure, still perform poorly compared to pre-defined molecular functions (*15*–*17*), which is also an effect of the lack of a quantitative classification of protein functionalities. Simultaneously, there exists no high-throughput screening system for emergent protein functions *in vitro*, putting an extra layer of difficulty to find promising candidates to eventually test in *in vivo* environments.

Here, we tackle these problems by combining ML-based distant homolog generation and subsequent functional screening of novel proteins for one of the best-studied model systems of biological self-organization, the bacterial Min system (*18*–*20*). The two proteins MinD and MinE evoke intracellular reaction-diffusion patterns resulting in ATP-dependent oscillations of the proteins between the cell poles in *Escherichia coli*, placing the division machinery at mid-cell and thus determining the division site (fig. S1). Remarkably, these spatiotemporal patterns, called Min waves, can be reconstituted *in vitro* (*21*), what has allowed the extensive biophysical characterization of the molecular dynamics of Min proteins, such as protein-protein and protein-lipid interactions that give rise to the emergent behavior. In particular, MinE has been dissected to a core set of molecular functions that are essential for spatiotemporal self-organization (*22, 23*), putting into reach the possibility to computationally predict these three molecular functions and hence to indirectly predict the emergent functionality. Thus, the emergent function of the MinDE system can be screened by combined *in silico, in vitro*, and *in vivo* (i³) setups (Fig. 1). In the following, we show that such an i^3^-screening pipeline can find novel, ML-generated homologs of MinE that can fully substitute the wildtype gene in a living organism, opening the door to targeted engineering of cellular functions by protein design.

**Fig. 1.**
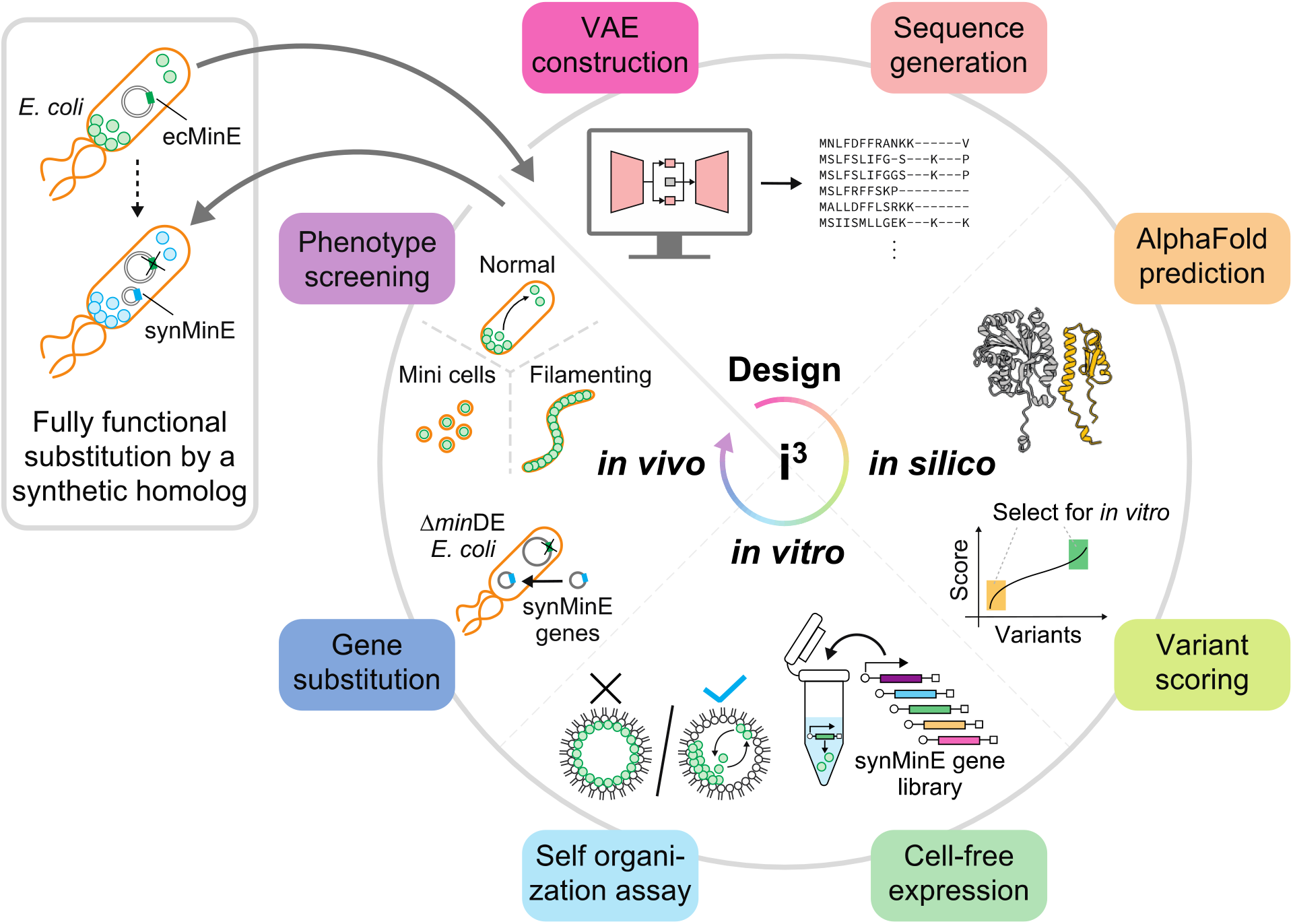
Overview of machine-learning assisted protein design and i^3^-screening pipeline for novel artificial homologs. The combined pipeline of this study: (1) Variational Auto Encoder-based sequence generation, (2) *in silico* scoring of synMinE structures as predicted by AlphaFold2, (3) *in vitro* self-organization assay for synMinE variants via cell-free expression, and (4) *in vivo* substitution of synMinE genes in ∆*min*DE *E. coli* cells. This pipeline finds an artificial, fully functional, distant homolog of the wildtype protein.

### Generation of artificial distant MinE homologs

To generate MinE-like proteins that we could subsequently screen for emergent function, we generated novel distant variants of wildtype MinE using a Multiple Sequence Alignment based Variational Autoencoder (MSA-VAE) as introduced by ref. *24* (Fig. 2A). We chose this architecture as it is one of the few methods that is experimentally validated and was shown to have a high success rate (*24*). Furthermore, there are relatively few naturally occurring MinE sequences to train the model on, compared to other studies where novel homologs were generated (∼6,000 sequences in our dataset, compared to ∼17,000 sequences used to train ProteinGAN, (*25*)), so we preferred the MSA-VAE over a GAN as it needs fewer parameters because information is already encoded in the MSA (ProteinGAN has ∼60,000,000 trainable parameters while our model has ∼1,000,000 (*25*)). We trained the MSA-VAE with a modified ELBO loss function similar to ref. *24* with a range of different hyperparameters (see Methods) and evaluated performance by single and pairwise amino acid frequency distributions as in the original paper (fig. S2). A high correlation in this metric indicates that evolutionary constraints are considered when generating sequences, rather than simply introducing random noise (*24*). With the selected set of hyperparameters, we generated 4000 novel variants. As can be seen in Fig. 2B, sequence conservation among the generated variants is highly similar to sequence conservation in naturally occurring variants, indicating that the model had generated reasonable sequences instead of just introducing random mutations. Dimensionality reduction on the latent space by Principal Component Analysis (fig. S3) further showed clustering by phylogenetic groups, confirming that the latent space conserved information about sequence relationships. However, some overlaps in the clusters can be observed and the correlation between pairwise amino acid frequencies of natural and generated sequences is not perfect, indicating that the generative model might still introduce some mutations that could impair the function of the protein. Thus, we had generated a set of artificial homologs where different grades of functional performance were expected, which we could then subsequently screen for emergent function, first *in silico* and then *in vitro*.

**Fig. 2.**
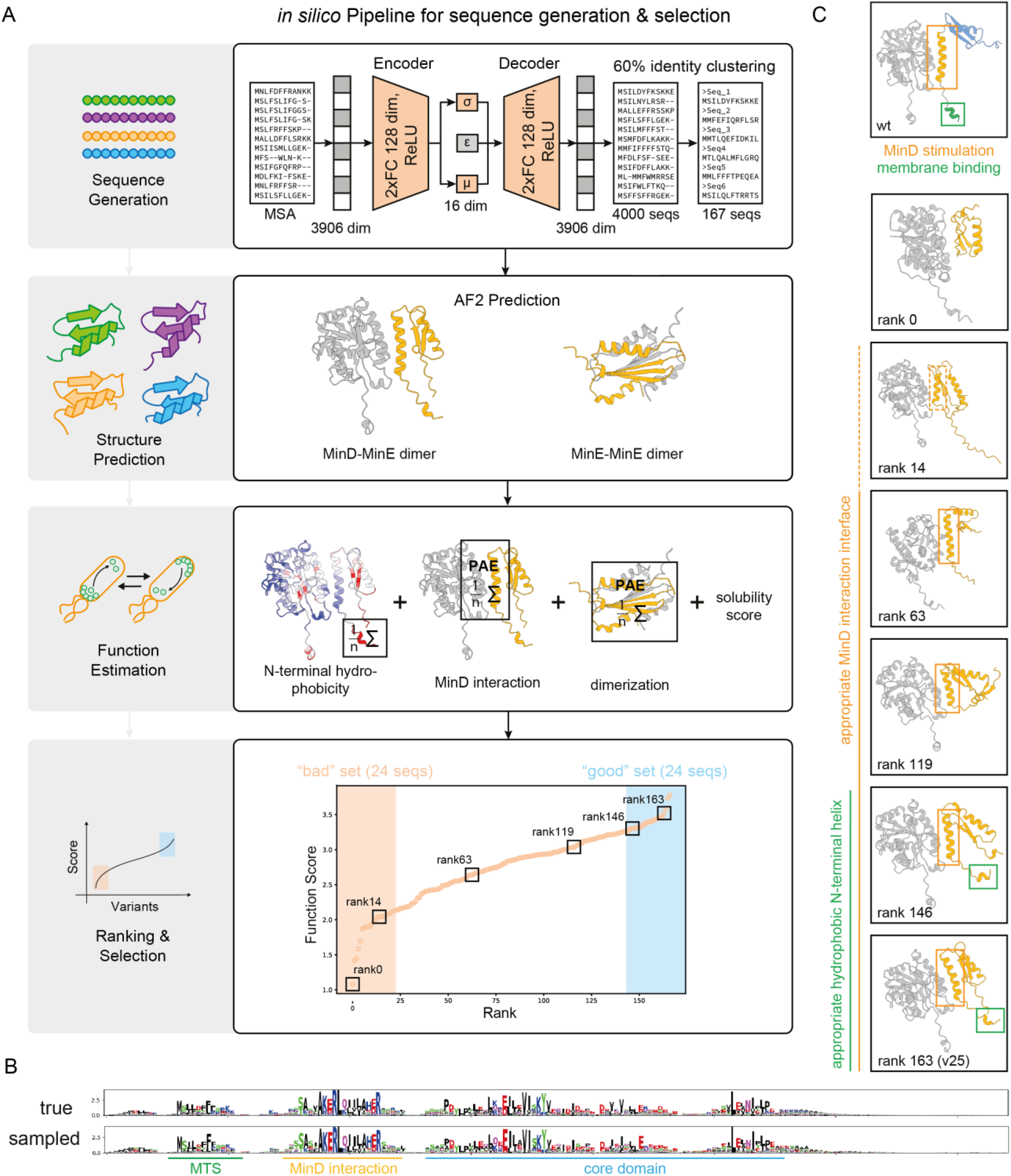
Generation of distant MinE homologs and *in silico* screening for expected function. **(A)** Pipeline overview. Sequences are generated using a Variational Autoencoder and clustered by 60% identity to ensure heterogeneity. The structures of the remaining 167 sequences are predicted using AlphaFold2 for homo- and heterodimers. A function score is calculated based on solubility and the three properties known to allow MinE to oscillate in *E. coli*: N-terminal membrane binding, interaction with MinD, and dimerization. The results are ranked, and the best and worst 24 candidates are selected for *in vitro* analysis. (**B**) Sequence conservation in naturally occurring and newly generated MinE homologs are highly similar. (**C**) Visual validation of ranked structures. With better ranking, structures show better MinD-interaction sites and membrane targeting regions.

### In silico scoring of emergent function

To reduce the number of proteins to be evaluated from 4000 to a more feasible number for subsequent screenings, and to ensure heterogeneity among the proteins, we initially screened proteins based on sequence identity. First, we filtered out all proteins with more than 60% sequence identity to the wildtype MinE in *E. coli*, as we eventually wanted to test the function in this organism. Second, we clustered the remaining generated variants by 60% sequence identity. Third, we randomly selected one sequence per cluster for further analysis. As a result, we got 167 remaining sequences to evaluate in our *in silico* pipeline.

The higher-order function of the MinE protein, volume oscillations through spatiotemporal self-organization with MinD, is known to be based on three properties (*20, 22, 23*): (i) membrane binding, (ii) formation of the MinDE complex that stimulates MinD’s ATPase activity, and (iii) homo-dimerization (Fig. 2C upper panel). To evaluate the expected functionality of the generated variants, we first predicted the structures of the variants using AlphaFold2 Multimer (*26*), and then developed a pipeline to estimate the three properties from the structure. Thus, we used a full sequence-structure-function pipeline (Fig. 2A). Interaction to MinD and dimerization was evaluated based on the Predicted Align Error (PAE) of the AlphaFold2 Multimer (*26*) output, similar to other protein design studies (*7, 8*). The membrane binding capability was estimated by calculating the hydrophobicity of the N-terminal region using ProteinSol Patches (*27*), since the hydrophobic interaction between the N or C-terminal region of proteins and lipid molecules is a determinant factor of the membrane binding (*28*–*30*) (see Methods and fig. S4A-C). As we eventually wanted to test the proteins in *E. coli* cells, we also predicted solubility of the proteins in *E. coli* using ProteinSol (*31*) as fourth score (fig. S4D). All four scores were normalized and summed up, resulting in a roughly normally distributed final “Function Score” (Fig. 2A lower panel). We then sorted the 167 heterogeneous variants by this score and validated the ranking visually. As can be seen representatively in Fig. 2C, proteins with low scores tend to be predicted to miss a proper interaction interface with MinD and to have a disordered and either very long or very short N-terminal region, suggesting impaired MinD’s ATPase stimulation and membrane binding. Proteins with high scores tend to resemble the wildtype closely. Interestingly, it is known that a conformational change is needed to swap from MinE-MinE homodimer to MinE-MinD heterodimer, and among the low-scoring variants such a change was often not predicted (fig. S5). As this could either be a truly missing property of the protein, in which the score would be correctly low, but also a misprediction of AlphaFold2 Multimer, in which case the score might be falsely low, we subsequently chose the best-scoring and worst-scoring 24 sequences for experimental screening (Fig. 2A lower panel, fig. S6, 7, and data file S1) and double-blinded them to be able to validate our *in silico* scoring approach.

### *In vitro* screening for emergent function using a cell-free system

Although the *in silico* screening effectively reduced the synMinE variants to test, it was not our intention to purify all 48 proteins at the same time, accompanied by many difficulties in experimental optimization due to protein solubility, cell toxicity, etc. Instead, to accelerate the screening pipeline, we utilized an *in vitro* cell-free protein synthesis system (*32*–*34*), where target proteins can typically be expressed within 1 h of incubation of a mixture of transcription-translational factors and DNA templates encoding the target proteins. Cell-free expression systems have been crucial in bottom-up synthetic biology, and have an enormous potential to be further utilized in various experimental setups by proper choice and optimization of the configuration, such as cell types and cell lysate or purified components-based systems. In this study, we chose the *E. coli*-based cell-free synthesis platform called PURE system (*35*), because it has been demonstrated that the PURE system can synthesize Min proteins, and that such cell-free expressed MinD and MinE proteins can self-organize into dynamic wave patterns *in vitro* in cell-mimicking environments such as lipid containers (*36, 37*).

We performed *in vitro* screening of the 48 novel MinE variants, named synMinEv1-48, by validating whether cell-free expressed synMinE variants could form Min waves in an established *in vitro* reconstitution setup. All 48 synMinE variants were synthesized with the PURE system (fig. S8), and subsequently, each variant was encapsulated within lipid droplets with purified MinD and ATP as cofactors. After checking for Min waves by laser scanning microscopy, we found in total 14 positive variants that give rise to spatiotemporal patterns on the lipid membrane with typical Min wave patterns (traveling wave and pole-to-pole oscillation), as well as typical oscillation periods (1-2 min) in lipid droplets as previously reported (Fig. 3A, B, fig. S9, and movie S1) (*36*). The other 34 variants did not show any heterogeneous localization over the usual time scale of Min oscillations (fig. S9, and movie S2). Intriguingly, but not surprisingly, after unblinding the variants, we found that 10 of these positive variants are from the high *in silico* scoring candidates and 4 variants are from the low *in silico* scoring candidates, suggesting that *in silico* screening for emergent function is possible by screening for the necessary molecular functions. This could be extended to a broad range of other emergent functions and should also be possible with entirely *de novo* designed proteins.

**Fig. 3.**
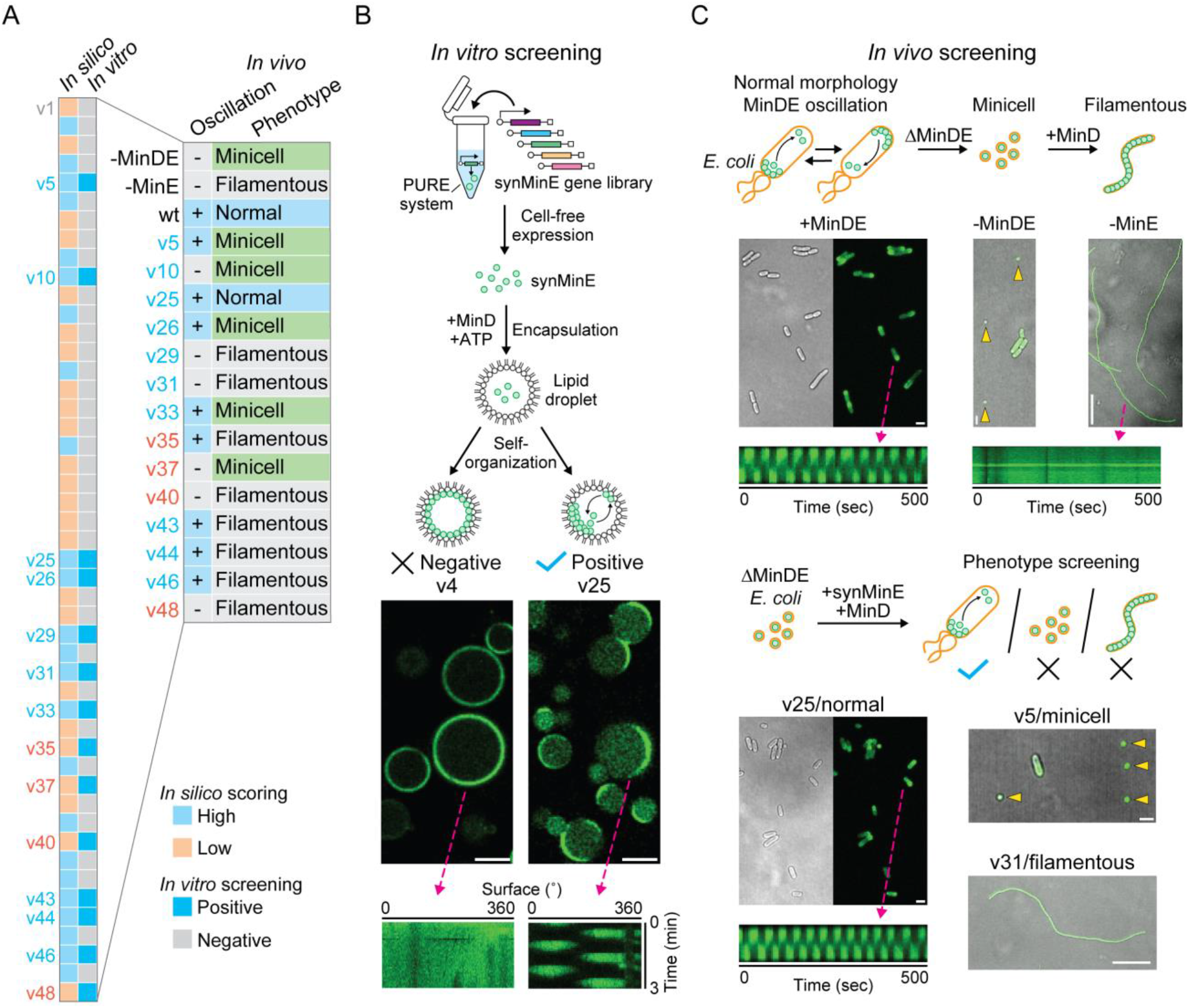
*In vitro* and *in vivo* screening for emergent functions of synMinE variants. (**A**) Summary of the i^3^-screening results for functional synMinE variants. The colors of each variant name indicate whether they are scored high (blue) or low (orange) by *in silico* screening. (**B**) *In vitro* screening of synMinE variants. Proteins were synthesized in the PURE cell-free expression system from a gene library of synMinE variants and then encapsulated in lipid droplets with EGFP-MinD (shown in green) and ATP. Emerging Min oscillation patterns as visualized in the kymograph on the bottom-right panel were observed with 14 variants from 48 candidates and further tested *in vivo*. Scale bars: 20 µm. (**C**) *In vivo* screening of synMinE variants. The remaining 14 variants were introduced in ∆*min*DE *E. coli* cells with GFP-tagged MinD. Subsequently, cell morphology and Min oscillation were validated to identify the functional synMinE variants. synMinEv25 fully substituted the wildtype (+MinDE condition). Differential interference contrast and fluorescence images are separately indicated for wildtype (+MinDE) and synMinEv25 (v25), or merged for the other conditions shown in the figure. Kymographs show Min oscillations inside the cells with wildtype or synMinEv25, while MinD did not induce oscillations without MinE (-MinE). Yellow arrows indicate the minicells in -MinDE and v5 conditions. Scale bars: 2 µm (+MinDE, -MinDE, v25, and v5) or 20 µm (-MinE and v31).

### *In vivo* substitution finds a fully functional complement of the wildtype gene

To further investigate whether these bottom-up constructed *in vitro* systems can truly screen for physiological function *in vivo*, we then assessed whether those 14 positive variants could also give rise to Min oscillations in *E. coli* cells. The 14 positive synMinE variants were introduced in an *E. coli* strain (HL1) lacking *min*DE genes by transforming plasmids encoding the respective synMinE variant and GFP-tagged MinD, as shown in previous studies (*23, 38*). With this setup, there are three possible phenotypes (*18*) (Fig. 3C, fig. S10, and S11): (i) the normal phenotype, (ii) the minicell phenotype, resulting from a lack or dysfunction of both MinD and MinE, where a certain number of cells (29% of the population in ∆*min*DE control vs 2.1% in the normal phenotype (i)) become non-chromosome miniature-sized spherical cells, due to the lack of the Min oscillation to place the division ring at mid-cell, and (iii) the filamentous phenotype, where MinE is lacking or dysfunctional but MinD is present and functional. This leads to homogeneous MinD binding to the cell membrane and thus, radical inhibition of the division ring assembly at any position. Indeed, in control setups of (ii: ∆*min*DE cells) and (iii: MinD expression in ∆*min*DE cells) without any transformed synMinE genes, Min oscillation can never be observed in cells (Fig. 3C).

Strikingly, we found that 7 out of 10 high-scoring synMinE variants evoked Min oscillations inside the cells, while only one of the low-scoring variants showed oscillations (Fig. 3A, C, fig. S10, S11, and movie S3-6). This suggests that the essential requirements for Min wave oscillation might be stricter *in vivo* than *in vitro*, potentially because proteins are constricted in smaller microscopic spaces and other cellular molecules, such as proteins, DNA, and RNA, could potentially induce non-specific interactions with the Min proteins. Furthermore, analysis of cell morphology revealed that the majority of wave-inducing synMinE variants, and especially all low-scoring variants, induce the minicell or filamentous phenotype, as a result of dysfunction in division ring assembly or placement (Fig. 3A, C, fig. S10, and S11). This suggests further complex molecular dynamics of Min proteins, as interaction of MinD with other proteins competing with MinE, such as the third Min protein, MinC (*20, 39, 40*), are essential to position the division machinery at the proper region. Finally, we found that one variant, synMinEv25, fully restores the normal cell phenotype together with Min oscillations (Fig. 3A, C, Fig. 4, movie S3, and S4), representing, to our knowledge, the first functional substitution of a natural gene by an artificial homolog generated by a generative model in a living organism. Additionally, these results also confirm that our *in silico* scoring could reasonably estimate the emergent protein function, and that, together with the *in vitro* screening, our pipeline considerably enhances the efficiency of the experimental validation for emergent functions.

**Fig. 4.**
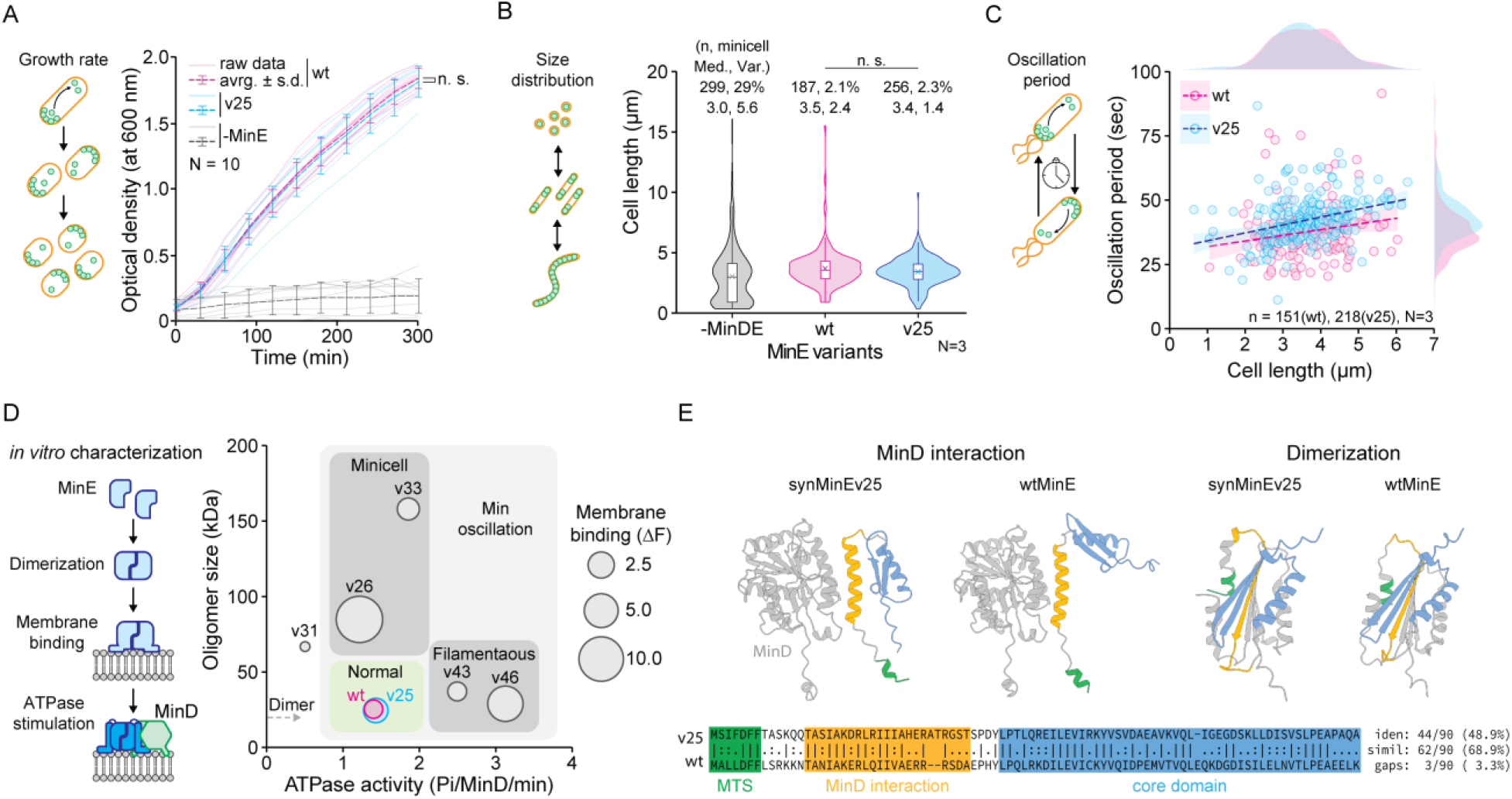
Characterization of synMinEv25 confirms the functional substitution of the wildtype. (**A**) Growth curves of *E. coli* cells show that the introduction of synMinEv25 in ∆*min*DE cells together with MinD recovers cell growth at the same level as wtMinE (n.s. indicates p > 0.05 between wildtype (wt) and synMinEv25 (v25) in Welch’s t-test at 300 min). Violin plots of cell-size distributions of *E. coli* cells confirm that both wtMinE and synMinEv25 confer proper size distribution while ∆*min*DE (-MinDE) cells produce a high population of minicells (< 1 µm in cell length). Box plots inside the violin distribution indicate maximum and minimum in 1.5xIQR, 25^th^ and 75^th^ percentile, median (bar), and mean (cross symbol) values. n.s. indicates p > 0.05 in Mann-Whitney U test. (**C**) Scatter plots of Min oscillations induced by wtMinE or synMinEv25 exhibit similar period and size dependency, confirming that synMinEv25 can functionally substitute the wildtype. The dotted lines and shades indicate linear trends with 95% confidence intervals. Distributions of plots are also shown as external density plots. (**D**) Bubble plot of *in vitro* characterization of synMinE variants. synMinEv25 has the closest scores to the wildtype among other variants, showing a fine match with the screening results. (**E**) Comparison between wtMinE and synMinEv25 structures and sequences confirms that synMinEv25 is a proper distant homolog while keeping similar structures to the wildtype.

### Functional analysis of synMinEv25 reveals its impeccable capability

To further understand the function of synMinEv25, we conducted *in vivo* and *in vitro* characterization of synMinE variants. First, we analyzed cell growth with all high-scoring synMinE variants. In contrast to the control (-MinE) condition, the introduction of synMinEv25 successfully restored growth rates to the wildtype-level (Fig. 4A). Also, 6 of the 10 positive high-scoring variants restored cell growth as well (fig. S12), suggesting that even without proper positioning of the division machinery inducing abnormal phenotypes, synMinE variants can induce cell division and growth. We then measured the cell size distribution of normal and minicell phenotype mutants to assess the accuracy of cell division led by Min oscillation. We found that synMinEv25 has a similar minicell population (2.1% (wt) vs 2.3% (v25)), median cell size (3.5 µm (wt) vs 3.4 µm (v25)), and even narrower size distribution than wildtype (2.4 µm (wt) vs 1.4 µm (v25) in variance), suggesting synMinEv25 allows proper cell division by placing the division ring in a correct location within comparable time- and geometrical-scales, and especially suppresses the production of elongated cells to confer better functionality in cell division (Fig. 4B). The other variants induced much higher minicell populations (7.2% ∼ 35%) and wider size distributions (fig. S13), meaning they were inefficient in positioning the cell division machinery. The Min oscillations induced by synMinEv25 indicated a similar tendency of periods against cell length as the wildtype, with slightly slower oscillations (Fig. 4C, fig. S14, movie S3, and S7). Taken together, synMinEv25 can substitute the wildtype in all intrinsic functions of the Min system - cell growth, morphology, and biological pattern formation.

Moreover, we set out to purify the promising synMinE variants and were able to obtain 6 out of the 10 high-scoring variants including synMinEv25 in a standard affinity purification protocol (fig. S15). This only 60% success rate further shows the great potential of cell-free expression for functional screenings. We characterized those purified proteins by three functional assays *in vitro* with regard to (i) membrane binding, (ii) catalyzing MinD’s ATPase activity, and (iii) oligomerization, where all three features are essential MinE functions and were estimated during the *in silico* scoring. Strikingly, a bubble plot (Fig. 4D, fig. S16, and S17) indicates that synMinEv25 has almost the same scores in all three parameters compared to the wildtype, showing that our screening and *in vivo* characterization are plausible. Moreover, we found an interesting relationship between scores and cell phenotype. Variants with higher ATPase induction activity than the wildtype but relatively similar oligomerization scores seem to induce the filamentous phenotype, while variants with ATPase induction comparable to the wildtype but bigger oligomer sizes seem to induce the minicell phenotype. This suggests that a delicate balance of those two parameters is particularly important for proper cell division, where the strength of membrane binding seems not to be a determining factor.

Finally, sequence comparison validated that the sequence identity of wildtype MinE and synMinEv25 is less than 50%, and sequence similarity is less than 70% (Fig. 4E), and, as can be seen in data file S1, sequence identity with its closest natural homolog is 78.7%. This shows that our generative model designed not just a mutant, but truly a distant homolog that can substitute a complex emergent function in a living organism.

Taken together, we have shown that our i^3^-screening system, including well-informed *in silico* scoring and cell-mimicking *in vitro* systems, can successfully screen for complex intracellular emergent functions. This marks a huge step forward towards customizable protein functions and the engineering of cellular functionality. Furthermore, we provided an example of a fully functional *in vivo* substitution of a gene found in nature by an ML-generated homolog, opening the door to the engineering of whole living organisms from the bottom-up.

## Acknowledgments

We would like to thank the MPIB Core Facility, Michaela Schaper, Katharina Nakel, Kerstin Röhrl, Beatrix Scheffer, and Sigrid Bauer for assistance in protein purification, plasmid cloning, *in vitro* characterization, preparation of competent cells, cell culturing, and lipid preparation. We are also grateful to Beatrice Ramm for her kind advice in HL1 cell preparation, to Karsten Borgwardt for useful feedback on the manuscript, and to Jürgen Cox for inspiring discussion. SK is supported by JSPS Overseas Research Fellowships. BF is supported by the Graduate School of Quantitative Biosciences Munich (QBM). LB is supported by the Max Planck School ‘Matter to life’.

