## Supplementary Materials for "Designing a protein with emergent function by combined *in silico, in vitro* and *in vivo* screening"

### Materials and Methods

Dataset construction

All sequences containing the InterPro (*41*) domain IPR005527 were downloaded from InterPro on 30/05/2022. They were clustered by cd-hit (*42*) with 100% identity cutoff, meaning that no redundant sequences were kept. As the dataset was small (8,496 non-identical sequences) and VAEs intrinsically add noise, no further clustering by identity was performed. All sequences shorter than 20 and longer than 200 amino acids and all sequences containing non-standard amino acids were omitted. A Multiple Sequence Alignment (MSA) was calculated using Clustal Omega (*43*), using the Hidden Markov Model Profile of the MinE domain provided by Pfam (*44*) (downloaded and extracted on 17/03/2022). To narrow the MSA, columns that contained gaps in over 98% of the sequences were cut out. The remaining dataset consisted of 5,958 sequences and the MSA was 186 columns wide. Finally, sequences were one-hot encoded as input for the VAE and split into a train (80%) and validation (20%) set. As the final evaluation of the model would be done experimentally, a test split was omitted.

Variational Autoencoder (VAE)

The Variational Autoencoder followed largely the architecture introduced by Hawinks-Hooker et al. (*24*) (Figure 2A), implemented in PyTorch (*45*). We optimized hyperparameters by evaluating the performance of the VAE using the correlation of pairwise amino acid frequencies (see below) of 4000 random samples generated by the VAE with the natural 5,958 sequences, a metric introduced by the original MSA-VAE paper (*24*). A high correlation in this metric indicates that evolutionary constraints are considered when generating sequences, rather than simply introducing random noise (*24*). In the final model, both Encoder and Decoder consisted of a fully connected neural network with two 128 dimensional hidden layers and ReLU activation function. The latent space was 16 dimensional. After the second hidden layer in the Decoder, a softmax function generates a probability score to observe an amino acid or gap at each position in the MSA. For optimization, the Adam optimizer was used with PyTorch defaults and batch size 8 and learning rate 0.001, and the model was trained for 60 epochs. As loss function a modified version of the ELBO loss was used, where the KL-divergence loss was multiplied with the factor 0.01. During hyperparameter optimization we had found that without such a weighting, the VAE would always generate the same sequence, similar to mode collapse in Generative Adversarial Networks (*46*). The final loss function used was

$$Loss = 0.01\,Loss_{KL} + Loss_{reconstruction} = 0.01\,KL\ +\ BCE =$$

$$0.01\frac{1}{D}\sum_{i=1}^{D}-0.5\left(1+\sigma_{log,i}-{\mu_i}^2-e^{\sigma_{log,i}}\right)-\frac{1}{L}\sum_{i=1}^{L}y_i\log(\hat{y}_i)+(1-y_i)\log(1-\hat{y}_i)$$

where KL is the Kullback-Leibler divergence between the latent distribution and a normal distribution, BCE is binary cross entropy, D is the number of dimensions of the latent space, L is the length of the one hot encoded sequence, $\sigma_{log,i}$ is the logarithmic variance of the i-th latent dimension, $\mu_i$ is the mean of the i-th latent dimension, $y_i$ is the true value of the i-th one hot encoding and $\hat{y}_i$is the predicted value of the i-th one hot encoding. $\sigma_{log,i}$ and $\mu_i$ are the output of the Encoder, $\hat{y}_i$ is the output of the Decoder.

VAE Evaluation

The sequence logo seen in Figure 2B was generated using the Python library logomaker (*47*). Amino acid frequency and pairwise amino acid frequency (fig. S2) were calculated with a custom python script, gaps were not taken into account. The projection on the latent space (fig. S3) was generated by encoding all 5,958 natural sequences in the latent space, calculating a Principal Component Analysis of the resulting 16-dimensional vector representations, and projecting on the first two principal Components. In the scatterplot in fig. S3 only the four largest phylogenetic groups are displayed for clarity.

Sequence Generation and Selection

With the final VAE model, 4,000 sequences were generated by randomly sampling from the latent space and decoding the 16-dimensional random vector to an amino acid sequence using the trained Decoder. As we wanted to make homologs distant to MinE in *E. coli* (ecMinE), we excluded all generated sequences that had >=60% identity with ecMinE (UniProt ID P0A734). Then, to ensure heterogeneity among the sequences to test, we clustered the remaining sequences by 60% identity using cd-hit (*42*) and randomly selected one sequence per cluster for further analysis. 167 sequences remained. All pairwise sequence alignments (Fig. 4E, data file S1) were calculated using browser version of the EMBOSS Needle pairwise sequence alignment tool (*48*) with default parameters.

*in silico* Function Estimation

The emergent function of MinE is known to be based on three properties (*20*, *22*, *23*): (i) membrane binding, which is mediated by a short hydrophobic N-terminal alpha helix, (ii) stimulation of MinD's ATPase activity by formation of a MinD-MinE heterodimer, for which a conformational switch in MinE, changing a beta-sheet to an alpha-helix, is needed, and (iii) the formation of homo-dimers. As we eventually wanted to score the generated sequences for the emergent function, we defined individual scores to estimate the capability to show each of the three individual properties, and then summed them up to a final "function score". As preparation, we used AlphaFold Multimer (*26*) to predict the structures of the generated novel MinE homologs under two conditions: first, in presence of *E. coli*'s MinD (UniProt ID P0AEZ3), thus testing for heterodimer capability, second, in presence of itself, thus testing for homodimer capability.

Membrane binding estimation

To evaluate the membrane binding capability, we calculated hydrophobicity with ProteinSol Patches (*27*), using the predicted heterodimer structure as input, as MinE must be able to bind the membrane while binding to MinD. We then calculated a single score by averaging the hydrophobicity score over all amino acids of the N-terminal alpha helix. If the N-terminal region was unstructured, we averaged over the full N-terminal region, stopping at the MinD interaction helix. fig. S4C shows a histogram of the resulting scores.

<u>MinD interaction estimation</u>

We evaluated the potential to interact with MinD based on the Predicted Align Error (PAE) matrix provided by AlphaFold Multimer. The PAE is a measure of confidence of AlphaFold Multimer (*26*), where small values indicate high confidence. Thus, if AlphaFold Multimer is confident about the interaction of two proteins, this value is low on average (*7*, *26*). The interaction of MinE and MinD is known to be located at a specific alpha-helix of MinE (*20*), and only this part of MinE is in a stable position relative to MinD, whilst the rest of MinE has some flexibility. Thus, we evaluated the potential to interact with MinD by averaging the PAE between the MinD-binding alpha helix of each generated MinE homolog and structured regions of MinD, while neglecting other parts of the MinE structure. fig. S4B shows a histogram of the resulting scores.

<u>Dimerization estimation</u>

We evaluated the potential to form homodimers similar to the MinD interaction, based on the PAE. We calculated the average PAE between structured regions of two novel MinEs, that is, alpha-helices and beta-sheets. As can be seen in fig. S4A and B, the PAE for dimerization was on average lower than for MinD interaction, indicating that most generated MinE homologs might dimerize, but not all of the might interact with MinD, or might interact correctly.

<u>Solubility prediction</u>

As we eventually wanted to test the novel homologs *in vivo* in *E. coli*, we used ProteinSol (*31*) to predict the solubility in *E. coli*. Values above 0.7 indicate a high probability of being soluble. As can be seen in fig. S4D, most homologs were predicted to be soluble.

<u>Final Scoring and selection for *in vitro* screening</u>

To merge the four individual scores (membrane binding, MinD interaction, Dimerization, Solubility) to one "function score", we normalized each score by setting the lowest value to 0 and the highest value to 1, and then summed them up, such that the final function score could reach values between 0 and 4. We then selected the highest scoring 24 and the lowest scoring 24 sequences for *in vitro* analysis (fig. 2A), double-blinded them, and named them synMinEv1-48 (data file S1).

<u>Preparation of synMinE gene library</u>

The amino acid sequences of each synMinE variant were reverse-translated into DNA sequences using the Codon optimization tool from Integrated DNA Technologies (Coralville, IA USA) to optimize codon usage by referencing *Escherichia coli* K12. Then, the sequence of the first 30 bp (10 amino acids) were further altered by maximizing the frequency of A and T bases while keeping the translated amino acids to optimize the cell-free expression yield. Then, 5' and 3' additional sequences (table S1) coding T7 promoter, Ribosome binding site, T7 terminator etc. were further attached to the synMinE sequences. The resultant 48 sequences were synthesized using eBlocks Gene Fragments service (Integrated DNA Technologies).

<u>Estimation of cell-free expression yield of synMinE variants</u>

Cell-free expression of synMinE variants was carried out using PUREfrex 2.0 (GeneFrontier, Chiba, Japan) according to the instruction from the supplier. Each synMinE gene was mixed in PURE solution at 1 ng/µL, together with 4.4 % of FluoroTect GreenLys *in vitro* Translation Labeling System (Promega, Madison, WI, USA) and then incubated for 4 h at 37 ˚C. Synthesized synMinE variants were then separated by sodium dodecyl sulphate poly-acrylamide gel electrophoresis (SDS-PAGE) and fluorescence was detected using Amersham Imager 600 (GE HealthCare, Chicago, IL, USA). The relative expression yield of each variant was analyzed using the Fiji software (*49*).

*In vitro* assay for functional synMinE variants

synMinE variants were synthesized using PUREfrex 2.0 as described in the previous section but incubating only 1 h at 37 ˚C without FluoroTect GreenLys *in vitro* Translation Labeling System. Then, expressed synMinE solutions were 5-folds diluted in the Reaction buffer (50 mM Tris-HCl, pH 7.5, 150 mM GluK, 5 mM GluMg) and further mixed with 1 µM EGFP-MinD, 2.5 mM ATP, and 10 g/L BSA to obtain the inner solution for the assay. The concentration of synMinE solutions was further varied up to 1- to 20-folds after checking the dynamics of MinD inside the lipid droplets at 5-folds dilution to confirm the emergence of Min waves at different concentration ranges of synMinE.

To prepare the lipid-oil mixture, 1-palmitoyl-2-oleoyl-glycero-3-phosphocholine (POPC) and 1-palmitoyl-2-oleoyl-sn-glycero-3-phospho-(1'-rac-glycerol) (POPG) (Avanti Polar Lipids, Alabaster, AL, USA) were mixed at 70:30 (POPC:POPG) mol% in chloroform at 25 g/L. Then, 50 µL of the POPC:POPG mixture was dried under nitrogen gas stream, and subsequently, 10 µL of decane (TCI Deutschland GmbH, Eschborn, Germany) and 500 µL of mineral oil (Carl Roth GmbH, Karlsruhe, Germany) were added to the lipid film, and lipids were resuspended in oil by vortexing for 1 min at room temperature. Then, 1 uL of the inner solution was added to 50 uL of the lipid-oil mixture and subsequently, emulsified by tapping to obtain lipid droplets.

For the observation of Min protein dynamics, 1 uL of the droplet solution was added in a well of a 384-well plate together with 50 uL of the lipid-oil mixture. Imaging of samples was carried out by a Zeiss LSM780 confocal laser scanning microscope using a Plan-Apochromat 20x/0.80 air objective (Carl Zeiss AG, Oberkochen, Germany), using a 488 nm Argon laser for excitation with 10 s intervals for 3-5 min to validate the self-organization dynamics of Min waves as previously reported (*36*). The visualization of images including kymographs were carried out using Fiji software and ImageJ macro published in ref. *37*.

Construction of plasmids encoding synMinE variants for *in vivo* observation and purification

All 14 positive variants found by the *in vitro* assay were cloned in a pMLB plasmid together with mGreenLantern-MinD as previously reported (*23*). Briefly, sfGFP was substituted by mGreenLantern (*50*) gene fragment (Integrated DNA Technologies) using GeneArt Seamless Cloning and Assembly Enzyme Mix (Thermo Fisher Scientific, Waltham, MA, USA) according to the supplier's protocol with sets of primers (table S1) from pMLB-sfGFP-MinD.MinE (*23*). Then, MinE (wildtype) was substituted with each synMinE variant using the Seamless Cloning method and primer sets (table S1). To construct pMLB-mGreenLantern-MinD and pMLB-mGreenLantern, MinE and MinD genes were omitted from pMLB-mGreenLantern-MinD.MinE plasmid using blunt end cloning technique. All enzymes for cloning (DpnI, T4 Phosphokinase, and T4 DNA Ligase) were purchased from Thermo Fisher Scientific. Additionally, all 10 positive variants with high scores of *in silico* screening were further cloned into a pET28 plasmid with C-terminus His-tag for purification. MinE (wildtype) was substituted with each synMinE variant using Seamless Cloning and primer sets (table S1) from pET28-MinE-His.

*In vivo* phenotype characterization, analysis of cell size distribution and oscillation period

Substitution of wildtype *min*E gene was carried out using *E. coli* ΔminDE strain, HL1 (Δ*min*DE *zcf*117::Tn10 *rec*A::*cat*). HL1 was transformed with the plasmids pMLB-mGreenLantern (as -MinDE condition), pMLB-mGreenLantern-MinD (as -MinE condition), pMLB-mGreenLantern-MinD.MinE (as wt condition), or pMLB-mGreenLantern-MinD.synMinEv5, as well as other 13 synMinE variants (as each variant condition). Transformed HL1 cells were inoculated from glycerol stocks and incubated in LB (with 100 µg/mL ampicillin) medium overnight at 37°C. Cells were then diluted to 1:100 in 50 mL LB (ampicillin) medium and grown at 37°C, 180 rpm for 90-180 min. After an optical density at 600 nm (OD600) of cell cultures reached ~0.1, Isopropyl-β-D-thiogalactopyranoside (IPTG) was added to the cultures at 50 µM to induce expression of Min proteins. Cells were further incubated at 37°C, 180 rpm for 2-3 h and then diluted in LB (ampicillin) medium with 50µM IPTG to an OD600 of 0.1.

To prepare agarose pads, 1% (w/v) of UltraPure Low Melting Point Agarose (Life Technologies, Carlsbad, CA, USA) was first melted in LB (ampicillin) medium with 50µM IPTG at 60 ˚C using a bench-top incubator. Then, 400 µL of agarose solution was pipetted onto a coverslip, and another coverslip was immediately placed on top of the agarose solution to obtain a planer surface of agarose pads. The agarose solution was left at room temperature for 30 min to obtain solid pads. The coverslip was removed from the top of the pad, and then cultured cells (1 µL) were spotted onto the agarose pad and left for 10 min at room temperature. Then, the agarose pad was flipped onto another coverslip and mounted to a Zeiss LSM780 confocal laser scanning microscope. Imaging was carried out as described for *in vitro* assay using C-Apochromat 40x/1.20 water-immersion objective (Carl Zeiss AG). The oscillation of Min proteins inside cells was captured with 5 sec intervals. The observation was performed within 2 h after sample preparation at room temperature.

The phenotypes of each synMinE variant were determined as (1) the filamentous: elongated (>25 µm in length) cells were observed, (2) the minicell: miniature size (<1 µm in length) cells were observed more than 6.3% of the population (which is three times higher than wildtype MinE (2.1%)), and (3) the normal. The phenotypes were confirmed to show the same morphology by at least three biological replicates. The cell size distribution and periods of Min oscillations were further analyzed by using Fiji software and custom ImageJ macro. Briefly, time-averaged fluorescence of mGreenLantern-MinD was used to determine cell position and length. Then, fluorescence was normalized along with a long axis of the cell to obtain 1-dimensional images and then vertically stacked at each time point, resulting in a kymograph. Then the period of Min oscillation was obtained by fitting the fluorescent signals (along with the vertical (time) axis) with a sine function.

Cell growth measurement

The *E. coli* HL1 cells transformed with pMLB plasmids were inoculated from glycerol stocks and incubated in LB (with 100 µg/mL ampicillin) medium for overnight at 37°C. Cells were then diluted to 1:100 in 50 mL LB (ampicillin) medium and incubated at 37°C, 180 rpm for 90 min (180 min in case -MinE and v31 conditions due to the slow cell growth) and the OD600 was measured as t = 0. Subsequently, 50 µM of IPTG was added to the cultures and cells were further incubated at 37°C, 180 rpm for 5 h. The OD600 of cell cultures were continuously monitored at each 30 min after addition of IPTG, and total 11 timepoints were measured per sample.

Purification of synMinE variants

Purification of EFGP-MinD, MinD, and MinE was described before (*23*, *51*), and the purification of all synMinE variants was performed based on wildtype MinE purification. In Brief, BL21(DE3) pLysS cells were transformed by pET28-synMinEv5 or the other variants, and then incubated in LB medium (with 50 μg/mL Kanamycin) for overnight at 37 °C. The overnight cultures were then diluted to 1:100 in 500 mL LB (Kanamycin) and incubated while shaking at 37 °C, 180 rpm. Then, IPTG was added at 1 mM to induce overexpression of synMinE variants once OD 600 nm reached 0.2-0.3. Cells were further cultured for 3-4 h and harvested.

The cell pellets were resuspended in Lysis buffer (50 mM Tris-HCl, pH 7.5, 300 mM NaCl, 10 mM Imidazole) and subsequently lysed using a tip sonicator (Branson ultrasonics S-250D, Thermo Fisher Scientific). The cell lysates were centrifuged for 30 min at 20,000 x g, 4 °C and then the supernatants were mixed with Ni-NTA agarose (QIAGEN, Hilden, Germany). The samples were then incubated for 10 min at 4 °C and loaded into a gravity column. Subsequently, Ni-NTA agarose was rinsed with Wash buffer (50 mM Tris-HCl, pH 7.5, 300 mM NaCl, 20 mM Imidazole, 10 % Glycerol), and then the proteins were eluted with Elution buffer (50 mM Tris-HCl, pH 7.5, 300 mM NaCl, 250 mM Imidazole, 10 % Glycerol). The buffer of the protein solution was dialyzed with Storage buffer (50 mM Tris-HCl, pH 7.5, 150 mM GluK, 5 mM GluMg, 10 % Glycerol) using Amicon Ultra-0.5 centrifugal filter unit 3 kDa (Merck KGaA, Darmstadt, Germany) and stored at -80 °C until further use. The concentration of the proteins was measured by Bradford Assay (Bio-Rad, Hercules, CA, USA), and separated by SDS-PAGE to check the purity.

Size exclusion chromatography

The oligomerization of synMine variants was estimated using ÄKTA pure with Superdex 75 Increase 10/300 GL column (Cytiva, Marlborough, MA, USA). The column was equilibrated with Reaction buffer prior to the measurements. The standard proteins (Blue Dextran2000: 2000 kDa, Aldolase: 158 kDa, Conalbumin: 75 kDa, Ovalbumin: 44 kDa, Carbonic Anhydrase: 29 kDa, RNaseA: 13.7 kDa, Aprotinin: 6.5 kDa) were separately loaded into the column and then eluted fractions were monitored to determine the peak fraction. The peak fraction of each standard protein was then fitted to obtain a standard curve by calculating

$$K_{av} = Ve - Vo/Vc - Vo$$

where $V_o$ is the column void volume (the elution volume of Blue Dextran2000), $V_e$ is the elution volume of each sample, and Vc is the geometric column volume (23.5 mL in case of Superdex 75 Increase 10/300 GL column).

The eluted fractions of each synMinE variant were monitored and then the elution volume was determined as the peak fraction. The oligomer size of synMinE variants was then calculated from the standard curve.

ATPase assay

ATPase assay was performed following the previous report using NADH-coupled assay (*23*). To prepare small unilamellar vesicles (SUVs), 1,2-dioleoyl-sn-glycero-3-phosphocholine (DOPC) and 1,2-dioleoyl-sn-glycero-3-phospho-(1'-rac-glycerol) (DOPG) (Avanti Polar Lipids) were mixed at 70:30 mol% in chloroform at 25 mg/mL. Lipids were then dried under nitrogen gas stream and then hydrated in Min buffer (25 mM Tris-HCl, pH 7.5, 150 mM KCl, 5 mM $MgCl_2$) at 4 mg/mL. Subsequently, the solution was vortexed to obtain multilamellar vesicles and the solution was further extruded through a membrane with 50 nm pore size to break down to the small unilamellar vesicles.

For the measurement of MinD's ATPase activity, 0.2 mg/mL of SUVs solution, 1 mM ATP, 2 mM phosphoenolpyruvate, 0.5 mM NADH, the mixture of pyruvate kinase (600-1000 U/mL) and lactate dehydrogenase (900-1400 U/mL) (Sigma-Aldrich, St. Louis, MO, USA), 2 µM MinD, and 2 µM of MinE or each synMinE variant were mixed in the Min buffer. Then, the decrease in absorption at 340 nm was measured in a 96-well plate using the Spark multimode microplate reader (TECAN, Männedorf, Switzerland). To calculate the ATPase activity, the linear parts of the measured values of the NADH absorption were fitted to gain a linear regression curve.

Quartz Crystal Microbalance with Dissipation monitoring (QCMD) measurements

QCMD measurements were carried out following the previous report (*23*). Prior to each measurement, silicon dioxide (SiO2)-coated quartz crystal sensors (Biolin Scientific, Gothenburg, Sweden) were treated with a 3:1 mixture of sulfuric acid and hydrogen peroxide (piranha-solution). Subsequently, sensors were rinsed with ultrapure water, dried under a stream of nitrogen, and mounted in the flow modules of the Qsense Analyzer (Biolin Scientific). After baseline stabilization, supported lipid bilayers (SLBs) formation was induced through constant injection (flow rate: 0.15mL/min) of a 1 mg/mL mixture of SUVs solution (prepared as described in the method section for ATPase assay), in the TK buffer (20 mM Tris-HCl pH 7.5, 150 mM KCl), spiked with 5 mM CaCl2. The formed SLBs were washed with TK buffer until no frequency change was observed. Then, 150 µl of the 5 µM of each synMinE variant in TK buffer was flown over the sensor at 0.15 ml/min and the change in frequency was monitored at overtone F9. The measured frequency was normalized by averaging the value of 5 consequent measurements at each time point, and then the change in frequency was determined by subtracting the maximum value (as baseline) from the minimum value (as dropped frequency).

Data analysis and statistics

All statistical tests were carried out using the R software (*52*). Welch's t-test was used for cell growth analysis as a standard unpaired t-test to avoid multiplicity issues. The Mann-Whitney U test was used for cell size distribution analysis since distributions were not normally distributed, and therefore a t-test would not have been suitable. The sample size (shown as n) and biological replicates (shown as N) are indicated in the corresponding figures.

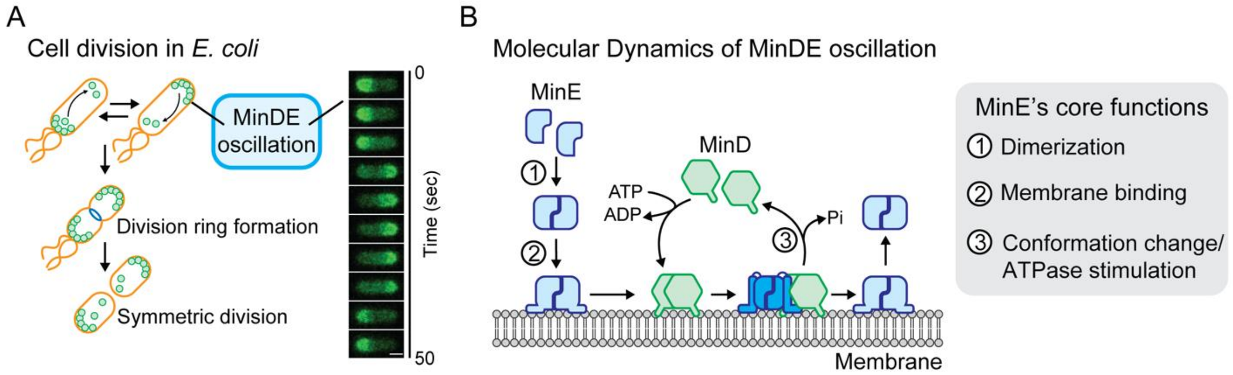


**Fig. S1. Schematic diagram of the MinDE oscillation.** (**A**) The oscillatory movement of MinD and MinE proteins determines the division site of *E. coli* cells at the mid-cell region. (**B**) Molecular dynamics of the MinDE oscillation. MinD is an ATPase that binds to the membrane upon ADP/ATP exchange. MinE forms a homodimer (1), binds to the membrane (2), and eventually forms the MinDE complex on the membrane (3) together with its conformation change to expose the MinD-interaction helix. Formation of the MinDE complex induces the ATP hydrolysis activity of MinD, and eventually ADP-state MinD and MinE detach from the membrane, forming periodical patterns on the membrane by repeating those processes.

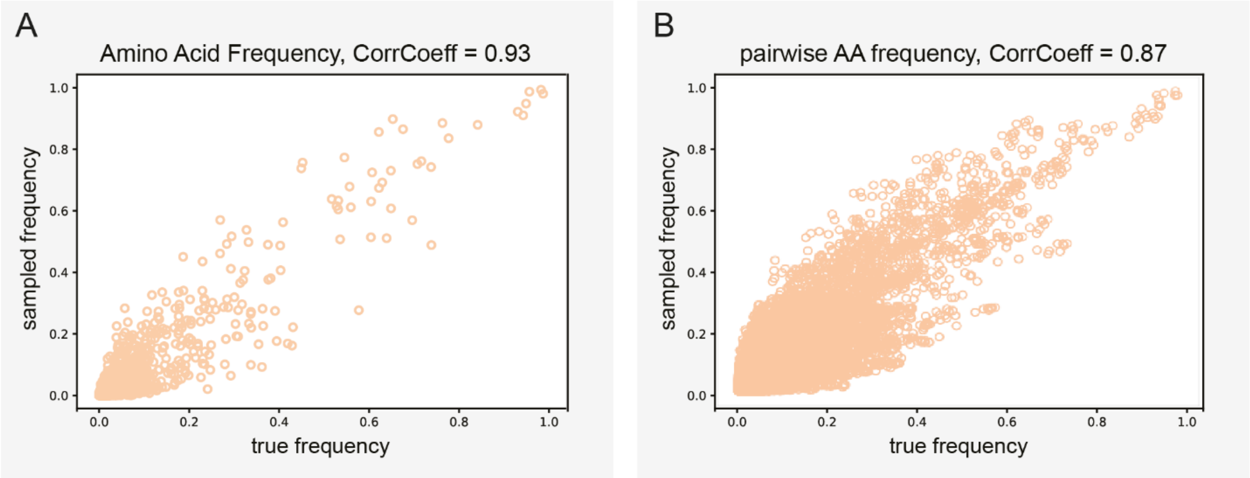


**Fig. S2. Evaluation of the VAE.** Scatterplot of amino acid frequency (**A**) and pairwise amino acid frequency (**B**) in natural MinE sequences (x-axis) and generated MinE sequences (y-axis).

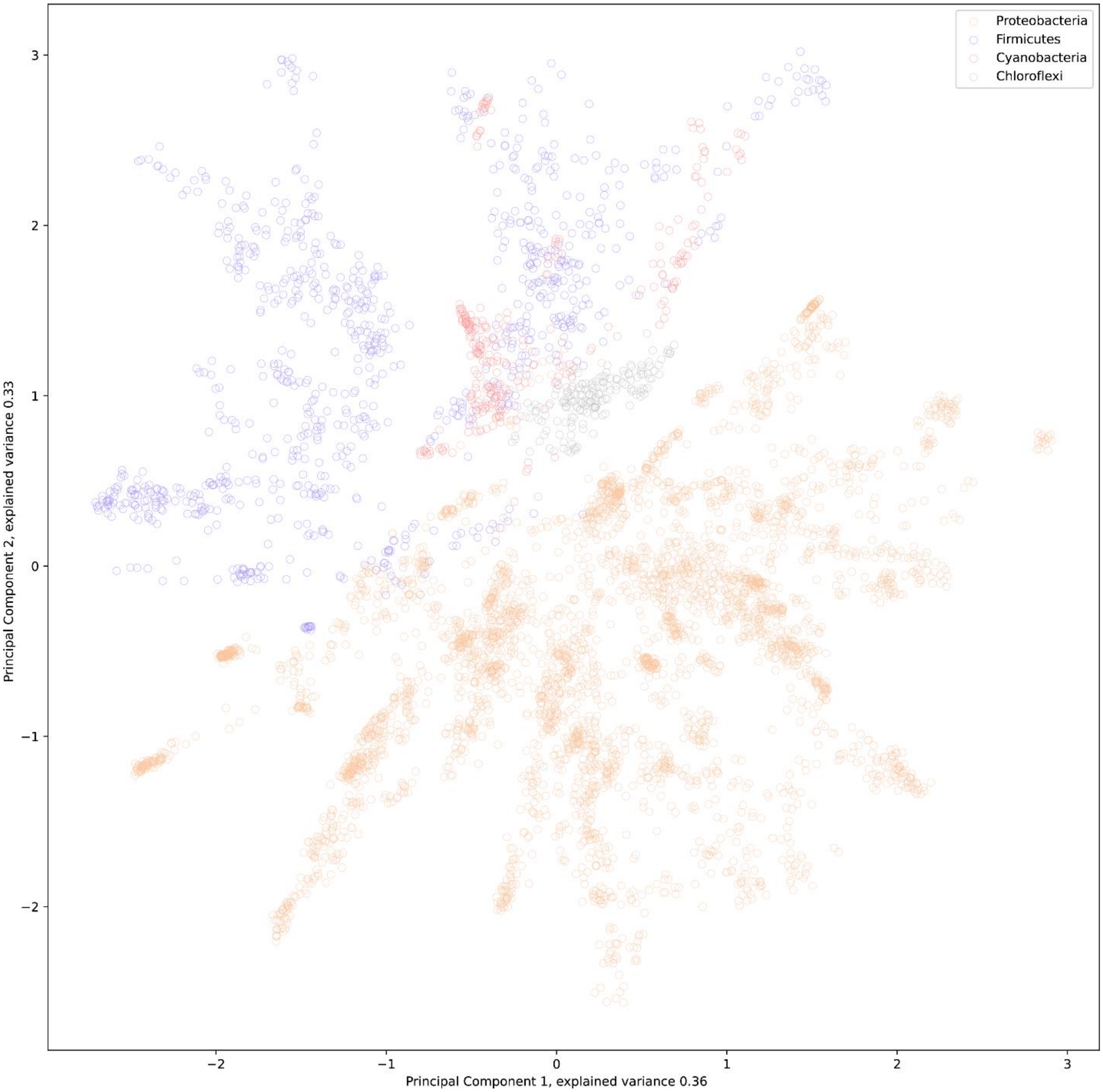


**Fig. S3. The latent space of the VAE conserves relationships among sequences.** Projection of natural MinE variants onto the first two Principal Components of the latent space. Each dot indicates a natural MinE and color indicates phylogenetic group. Only sequences belonging to the four largest phylogenetic groups are displayed.

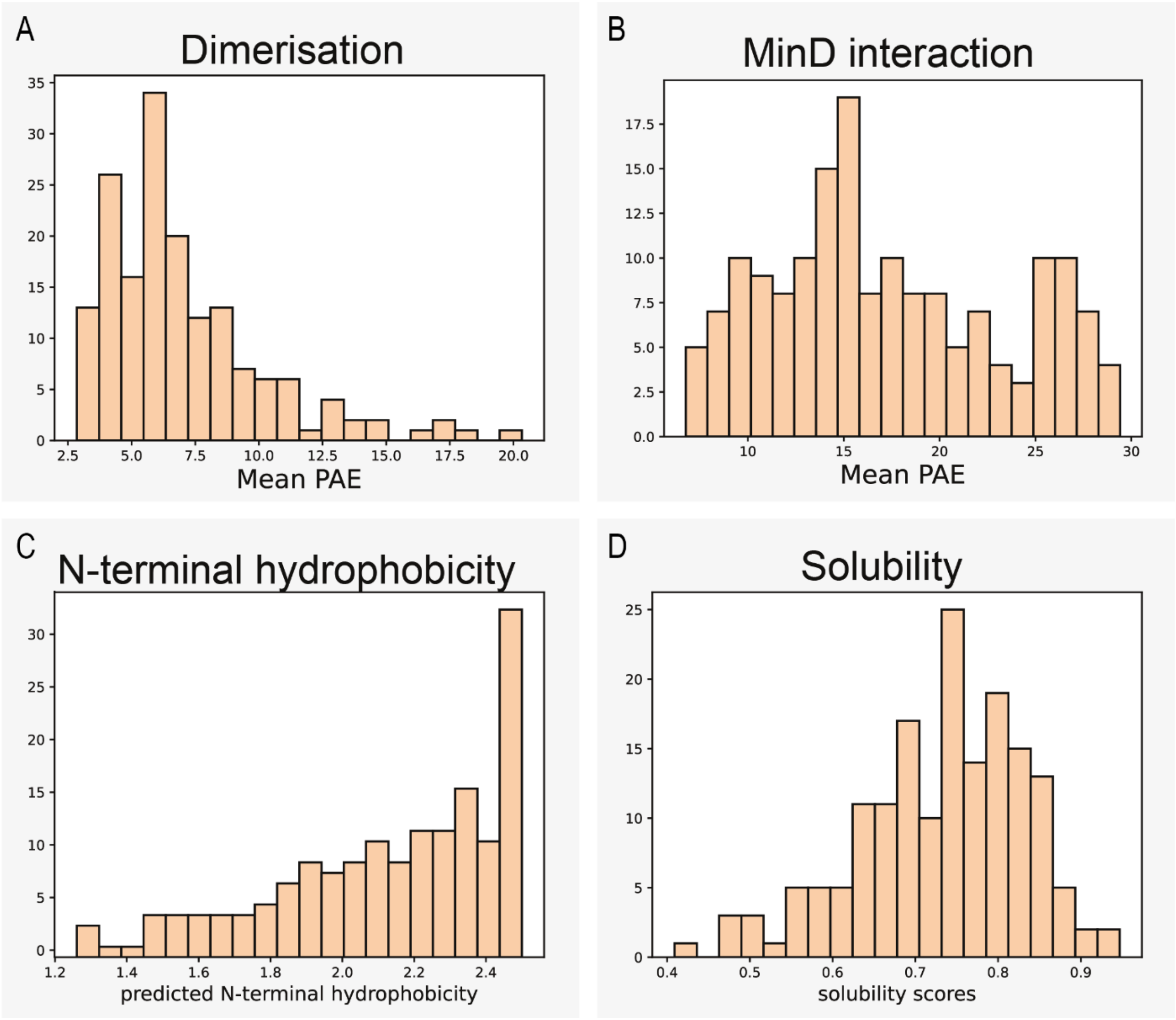


**Fig. S4. Histograms of individual scores of generated MinE properties.** (**A**) Dimerization scores, measured as average Predicted Align Error (PAE) between structured regions of two identical generated MinE proteins. Low scores indicate confidence about MinE dimerization. (**B**) MinD interaction score, measured as average PAE between a generated MinE's MinD-interaction helix and structured regions of MinD. Low scores indicate confidence about MinE-MinD binding. (**C**) Membrane binding scores, measured as average hydrophobicity predicted by ProteinSol Patches (*27*) at the N-terminal alpha helix. High values indicate high hydrophobicity and hence high probability of membrane binding. (**D**) Solubility scores, as calculated by ProteinSol (*31*). Values above 0.7 indicate a good predicted solubility.

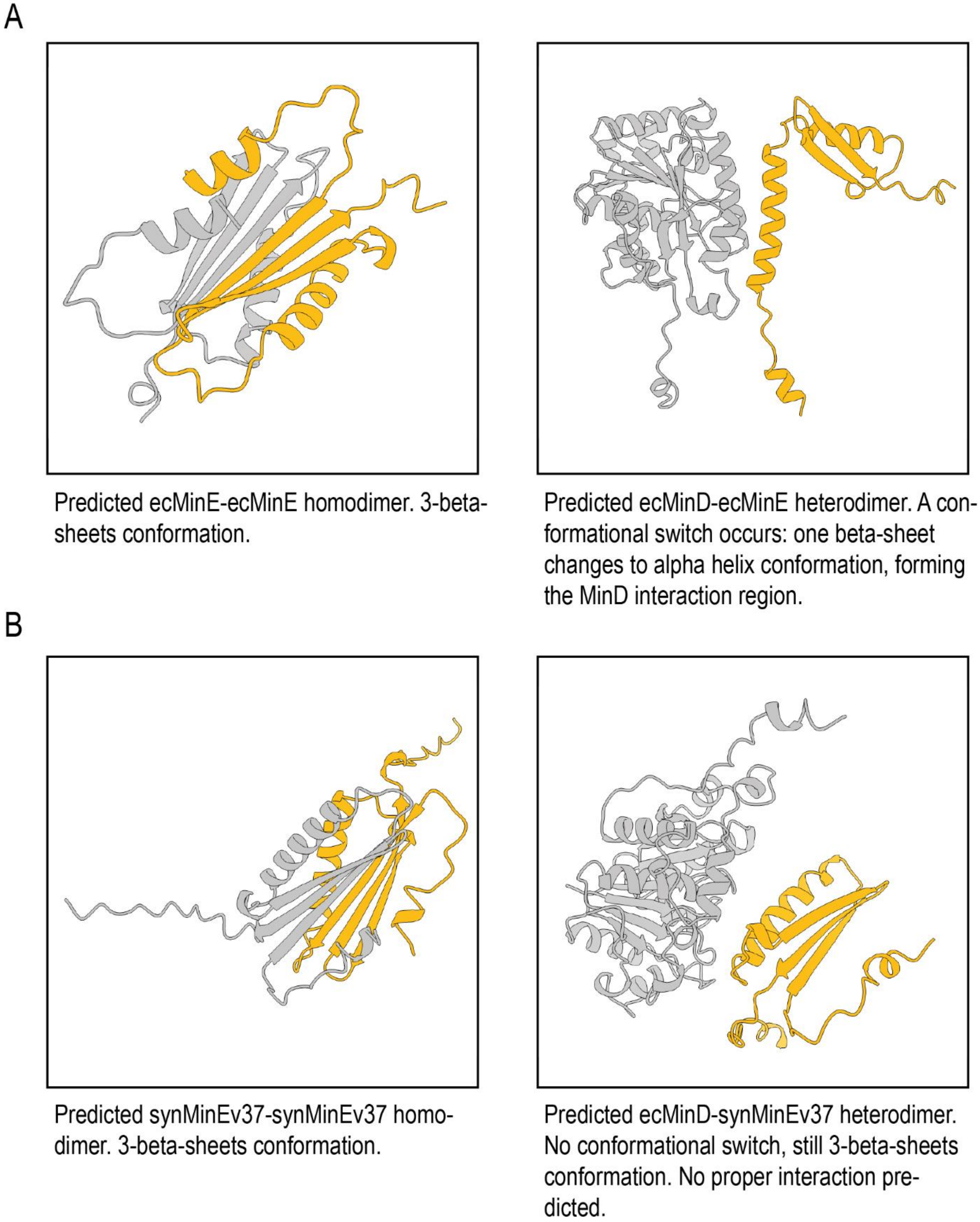


**Fig. S5. Missing conformational change in some low-scoring variants**. (**A**) wildtype, (**B**) synMinEv37. Especially, this occurred in three (synMinEv35, 37, and 48) out of the four variants that had low *in silico* function scores but did show oscillations *in vitro*, indicating that the score was falsely low due to misprediction of AlphaFold.

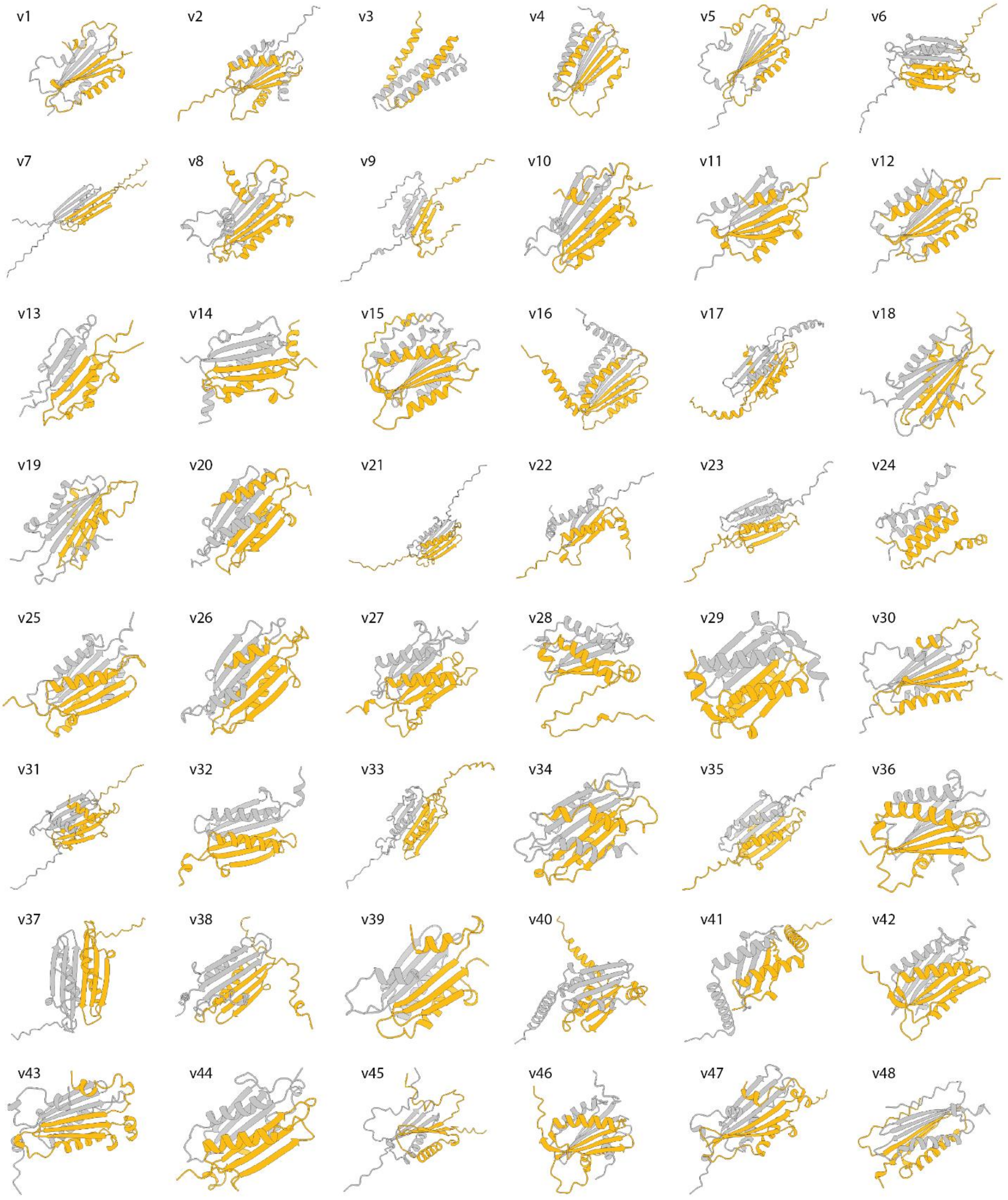


**Fig. S6. Homodimer structures of synMinEv1-48 as predicted by AlphaFold Multimer (*26*).**

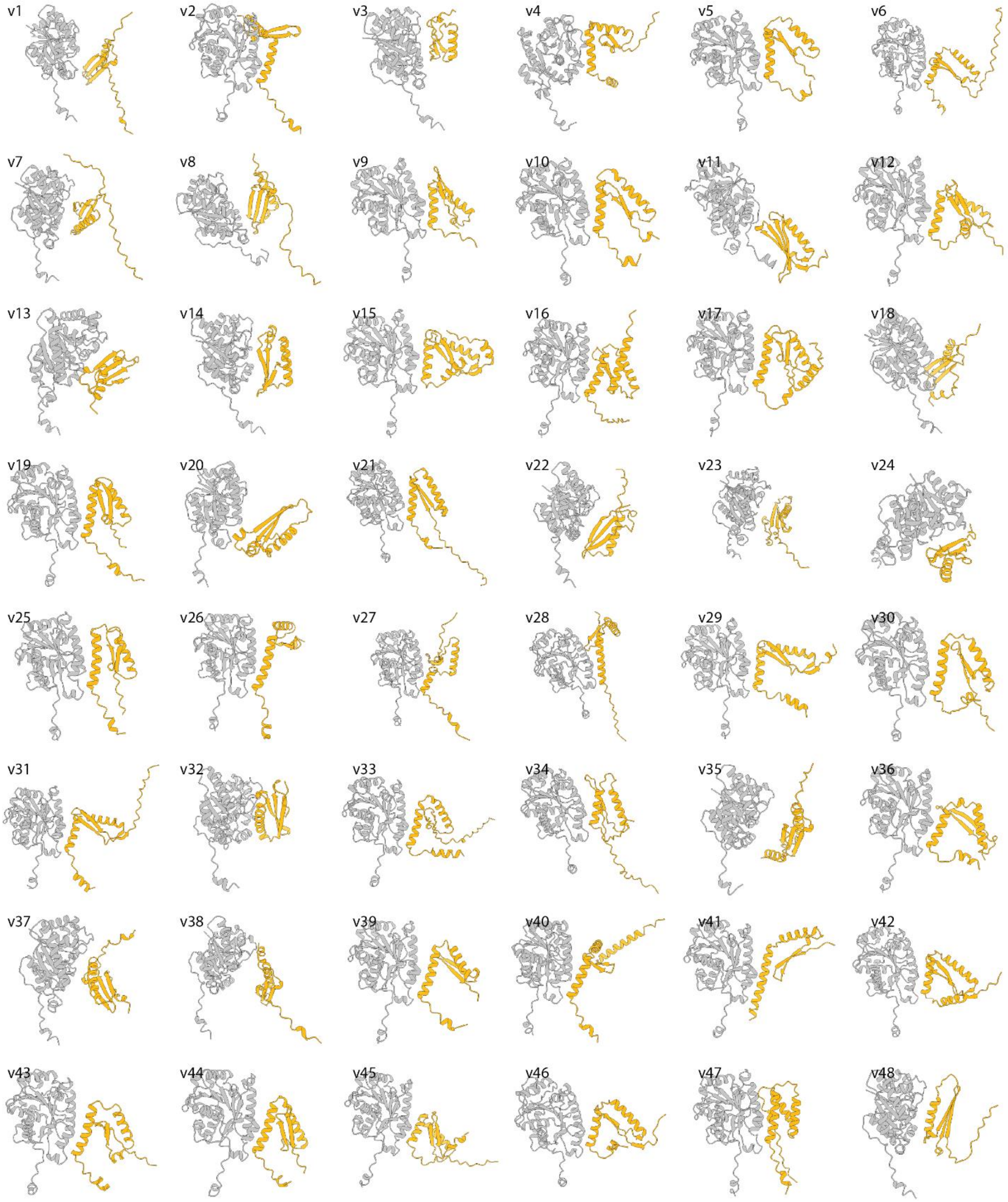


**Fig. S7. Heterodimer structures of synMinEv1-48 as predicted by AlphaFold Multimer (*26*).** ecMinD in grey, synMinE variants in orange.

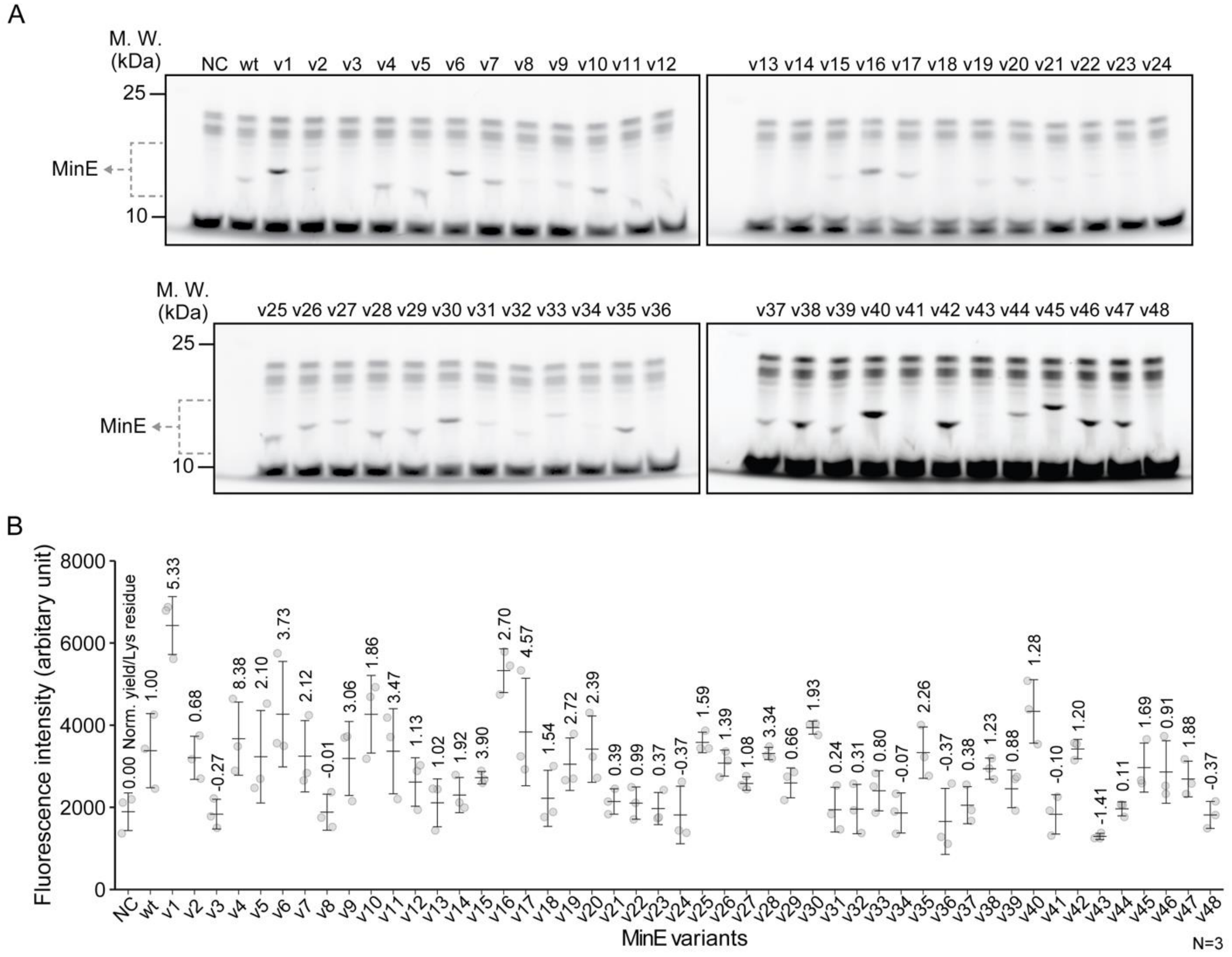


**Fig. S8. *In vitro* expression of synMinE variants.** (**A**) All synMinE variants were synthesized in the PURE cell-free expression system and detected by SDS-PAGE using the FluoroTect GreenLys *in vitro* Translation Labeling System. (**B**) Estimation of the cell-free expressed yield of synMinE variants confirms that more than 80% (40 variants) of synMinEs were synthesized in the PURE system at detectable levels, while 8 variants (V3, 8, 24, 34, 36, 41, 43, 48) did not get high yield by cell-free expression (Note: However, two of low-yielded variants (v43 and v48) were later positive in the *in vitro* screening (fig. S9), indicating such low yield of proteins are still sufficient to induce MinDE dynamics). Plots and bars indicate raw data, average, and standard deviation.

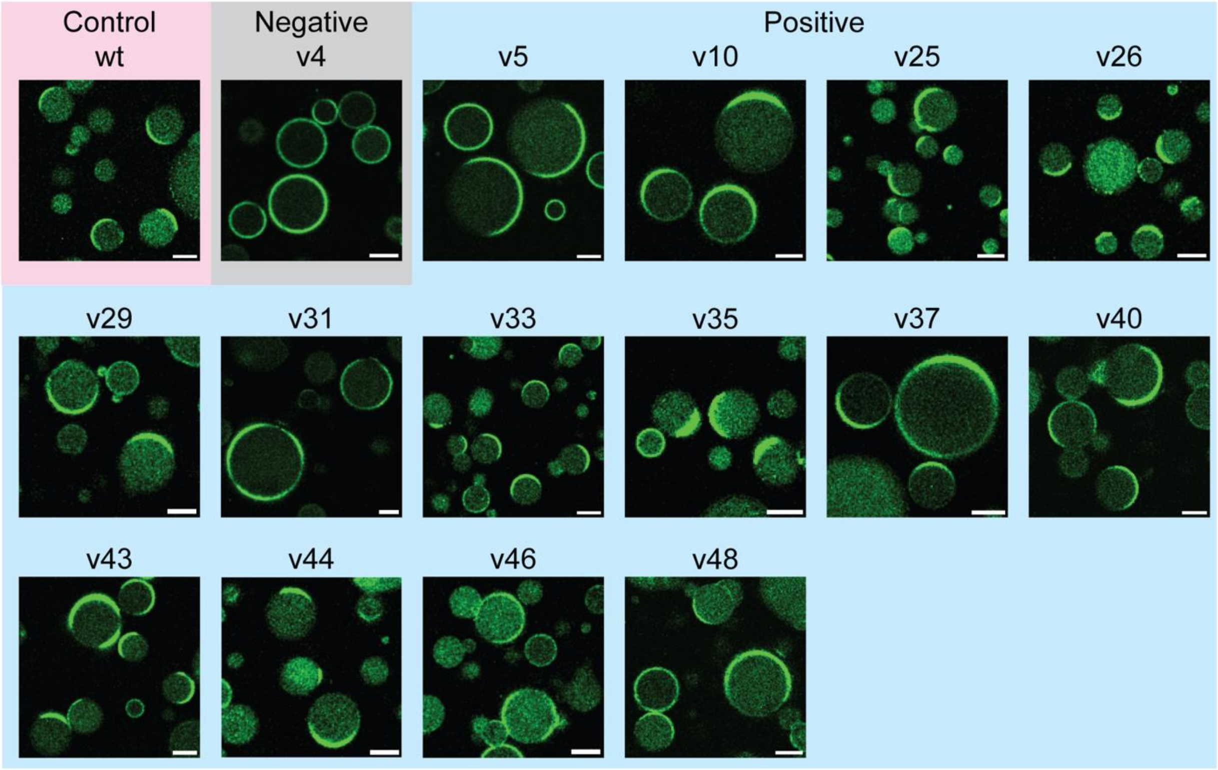


**Fig. S9.** ***In vitro*** **screening for functional synMinE variants.** All synMinE variants synthesized by the cell-free expression system were encapsulated in lipid microdroplets together with MinD and ATP, showing spatiotemporal pattern formation on the membrane with 14 "positive" variants, while the rest of the "negative" variants (v4 is shown as a representative example) did not induce dynamic behavior. Later, 10 of the 14 positive variants (v5, 10, 25, 26, 29, 31, 33, 41, 43, 44, and 46) were found to be high-scoring variants from the *in silico* screening.

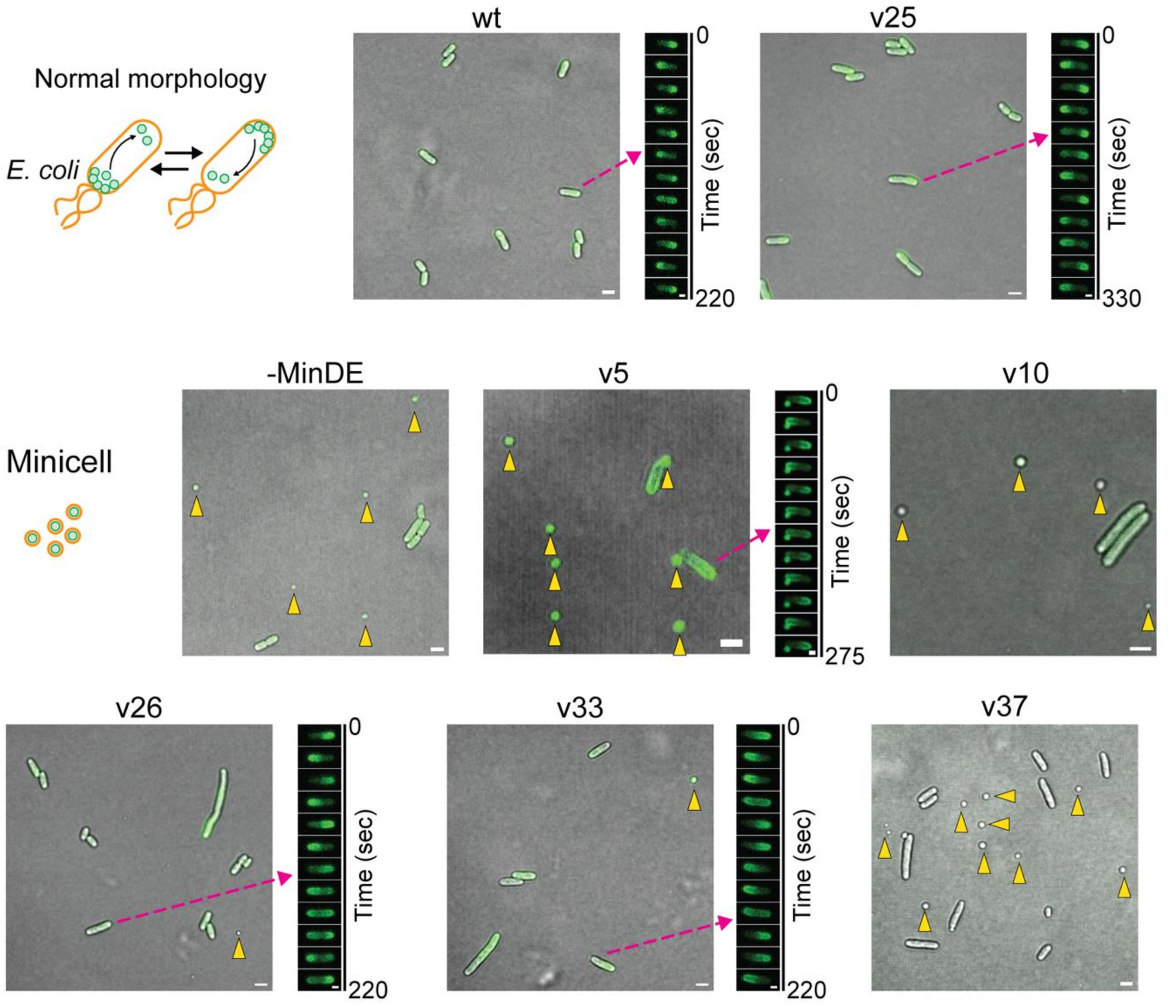


**Fig. S10. Normal and minicell morphology of *E. coli* cells induced by synMinE variants.** Wildtype MinE and synMinEv25 indicate normal morphology in Δ*min*DE *E. coli* cells with GFP-tagged MinD, showing that synMinEv25 fully substitutes the wildtype *in vivo*. 5 synMinE variants together with Δ*min*DE cells (-MinDE) induced minicells (indicated by yellow arrows) due to the lack of proper regulation of cell division, although some variants induced Min oscillations inside the cells. Differential interference contrast and fluorescence images are merged for a wide view of each condition, and fluorescence images are shown for visualizing Min oscillations. Scale bars: 2 µm for merged images (wide view) and 1 µm for the fluorescence images showing Min oscillations.

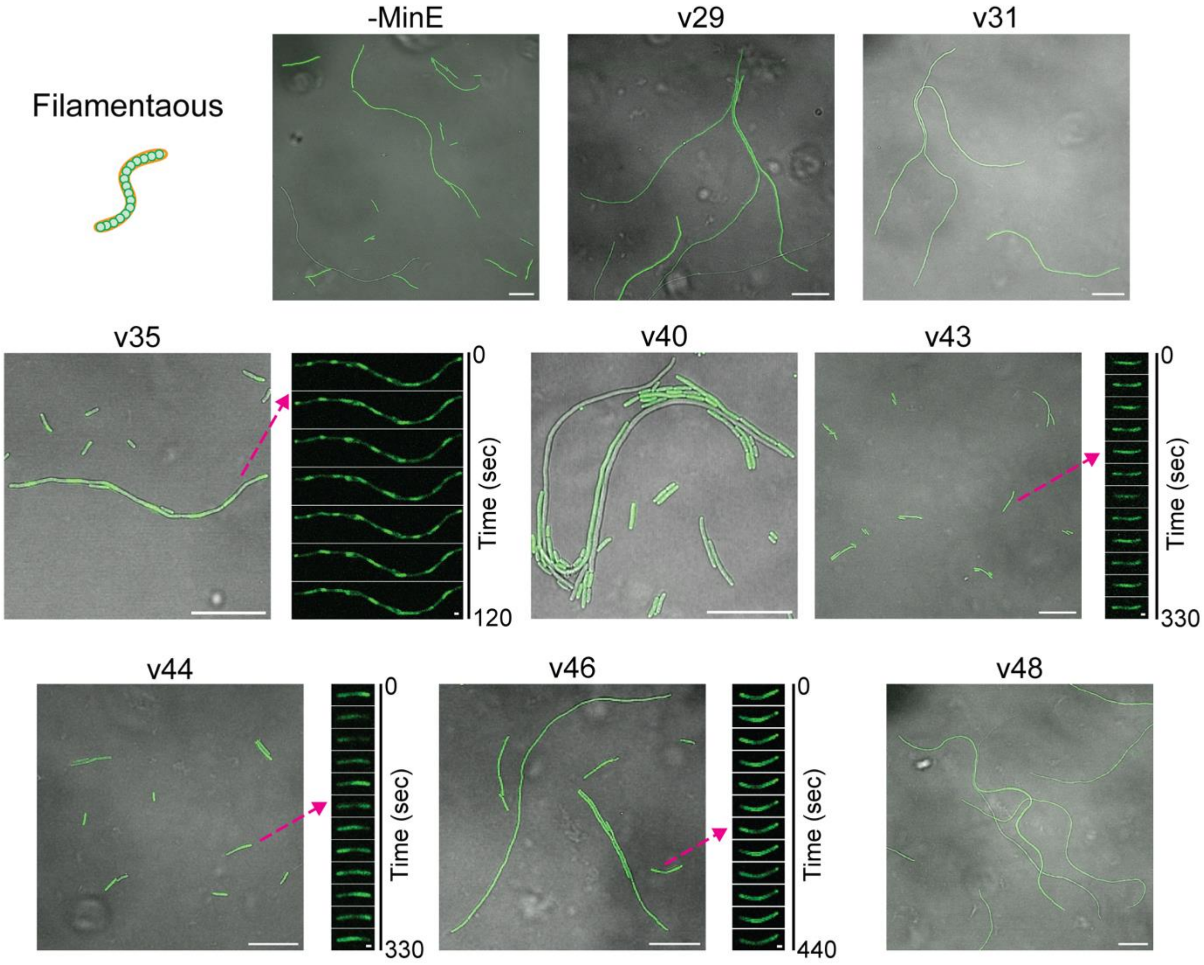


**Fig. S11. Filamentous morphology of *E. coli* cells induced by synMinE variants.** 8 synMinE variants together with Δ*min*DE cells transformed only with the MinD gene (-MinE) induced filamentous cells due to the inhibition of cell division. However, 4 variants indicate Min oscillations inside the cells, showing MinDE oscillations are partially functional in those conditions. Differential interference contrast and fluorescence images are merged for a wide view of each condition, and fluorescence images are shown for visualizing Min oscillations. Scale bars: 20 μm for merged images (wide view) and 2 μm for the fluorescence images showing Min oscillations.

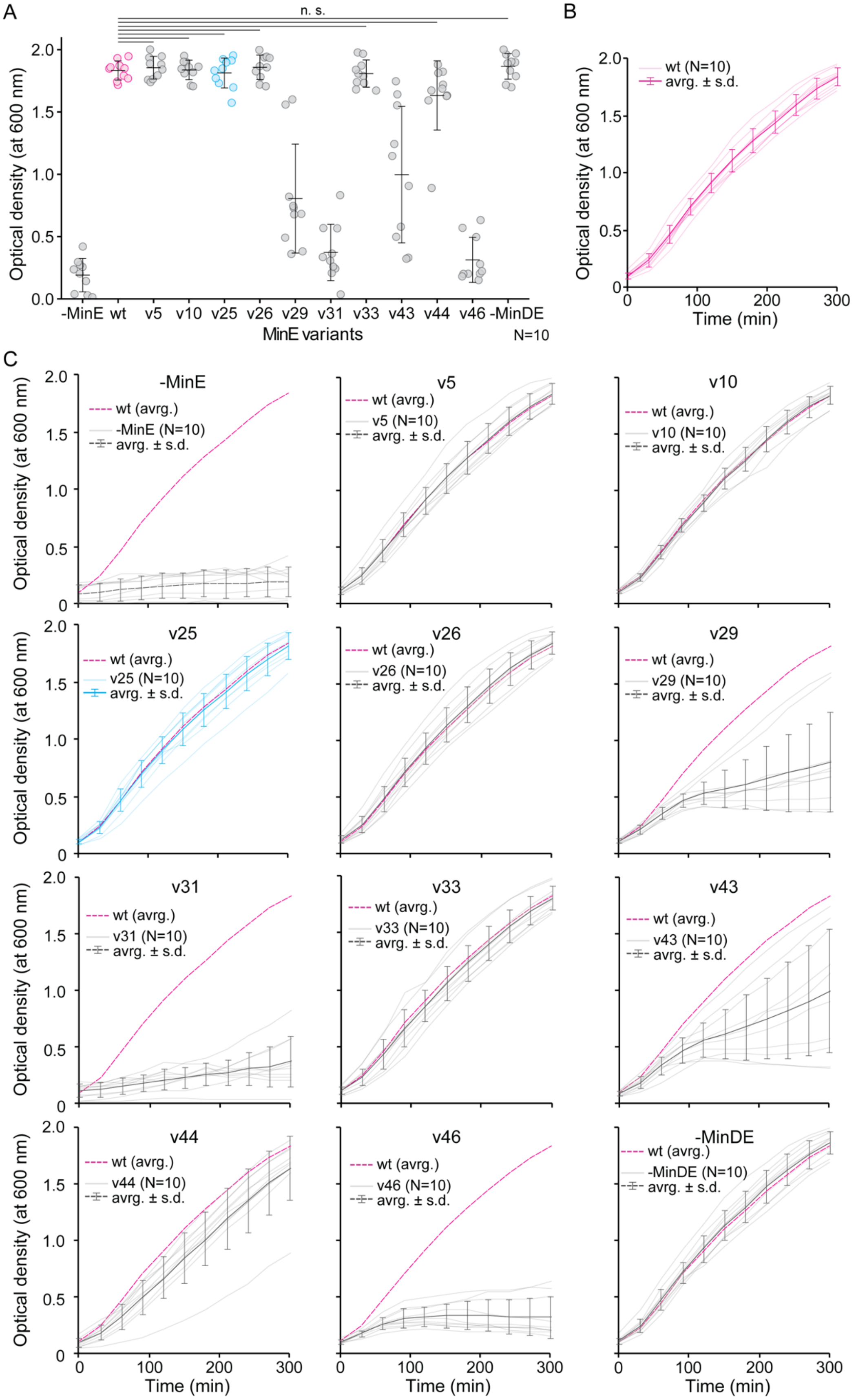
A
n. s.
Optical density (at 600 nm)
-MinE wt v5 v10 v25 v26 v29 v31 v33 v43 v44 v46 -MinDE
MinE variants
N=10
B
wt (N=10)
avrg. ± s.d.
Time (min)
C
-MinE
v5
v10
v25
v26
v29
v31
v33
v43
v44
v46
-MinDE
wt (avrg.)
-MinE (N=10)
v5 (N=10)
v10 (N=10)
v25 (N=10)
v26 (N=10)
v29 (N=10)
v31 (N=10)
v33 (N=10)
v43 (N=10)
v44 (N=10)
v46 (N=10)
-MinDE (N=10)
avrg. ± s.d.
Time (min)

**Fig. S12. Cell growth of *E. coli* cells containing synMinE variants.** (**A**) Growth of HL1 (Δ*min*DE) cells transformed with synMinE variants together with MinD (OD600 at 300 min of incubation) shows that 6 of 10 variants (v5, 10, 25, 26, 33, and 44) recovered cell growth at the same level as wildtype (n.s. indicates $p > 0.05$ between wildtype and synMinE variants in Welch's t-test). Plots and bars indicate raw data, average, and standard deviation. (**B**) The growth curve of the HL1 cell transformed with wildtype MinE together with MinD. (**C**) The growth curve of the HL1 cells transformed with synMinE variants together with MinD, only MinD without MinE (shown as -MinE), or none of them (shown as -MinDE) as a control.

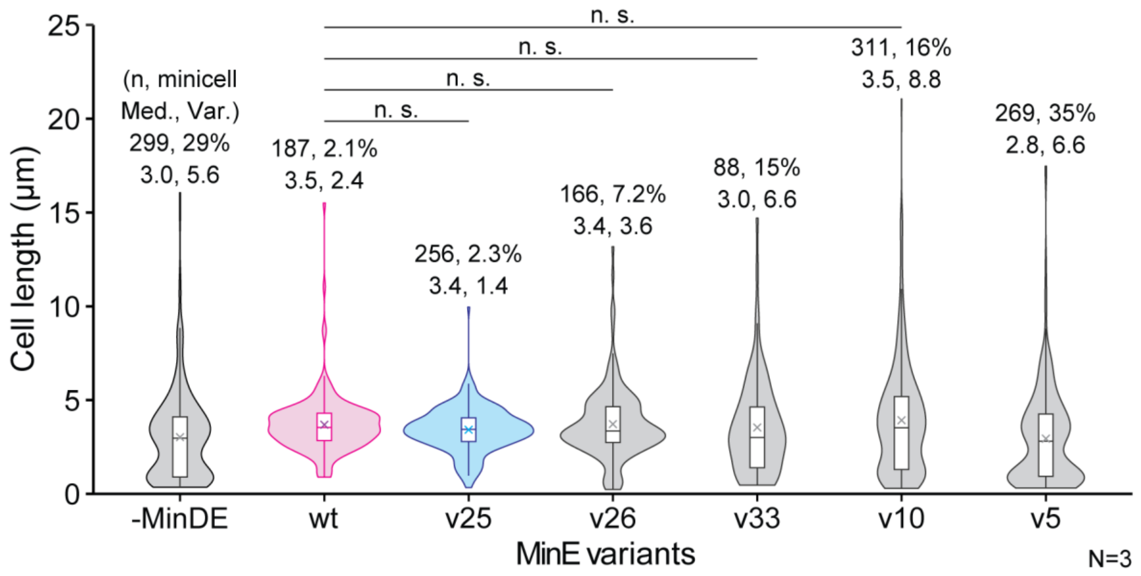


**Fig. S13.Violin plots of the size distribution of *E. coli* cells containing synMinE variants.** The size distribution of normal and minicell phenotype variants shows that synMinEv25 confers a similar size distribution to wtMinE, while Δ*min*DE (-MinDE) cells and other variants produce higher population of minicells (< 1 μm in cell length). Box plots inside the violin distribution indicate maximum and minimum in 1.5xIQR, 25th and 75th percentile, median (bar), and mean (cross symbol) values. n.s. indicates $p > 0.05$ in Mann-Whitney U test.

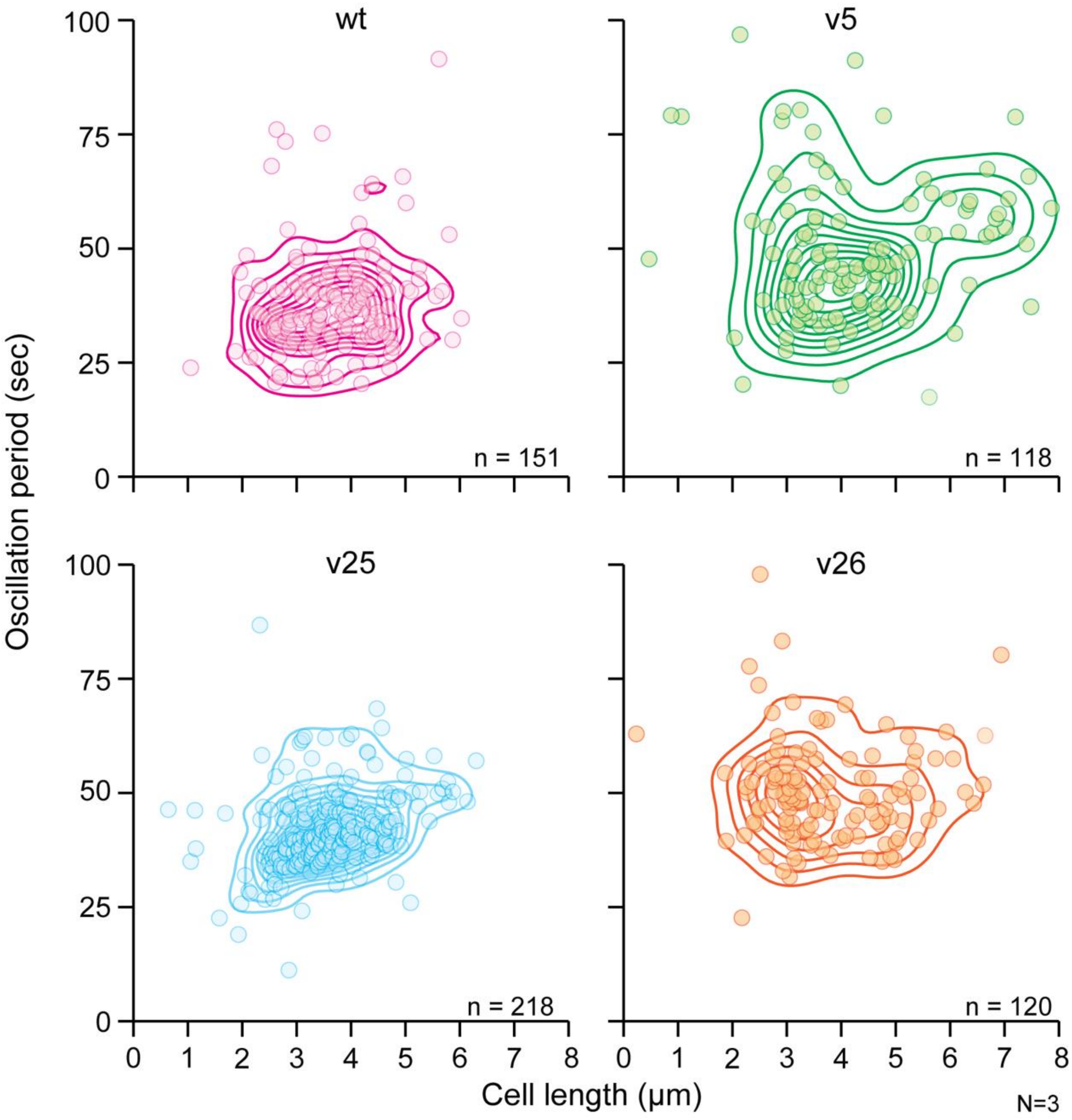


**Fig. S14. Scatter and density plots of oscillation period induced by synMinE variants.** Min oscillations induced by wtMinE or three synMinE variants, v5, 25, and v26 show similar period and size distribution in *E. coli* cells, indicating that synMinE variants properly function in *in vivo* environments.

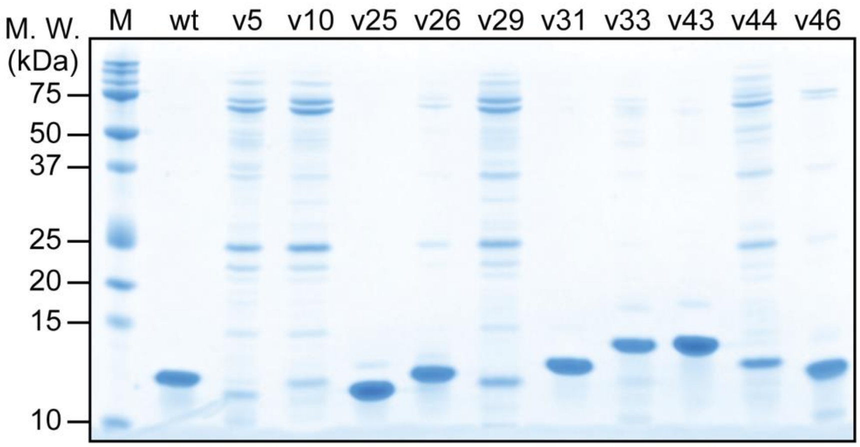


**Fig. S15. Purification of synMinE variants.** 6 of 10 synMinE variants (v25, 26, 31, 33, 43, 46) were obtained from a standard His-tag purification protocol at high yield and therefore used for *in vitro* characterization.

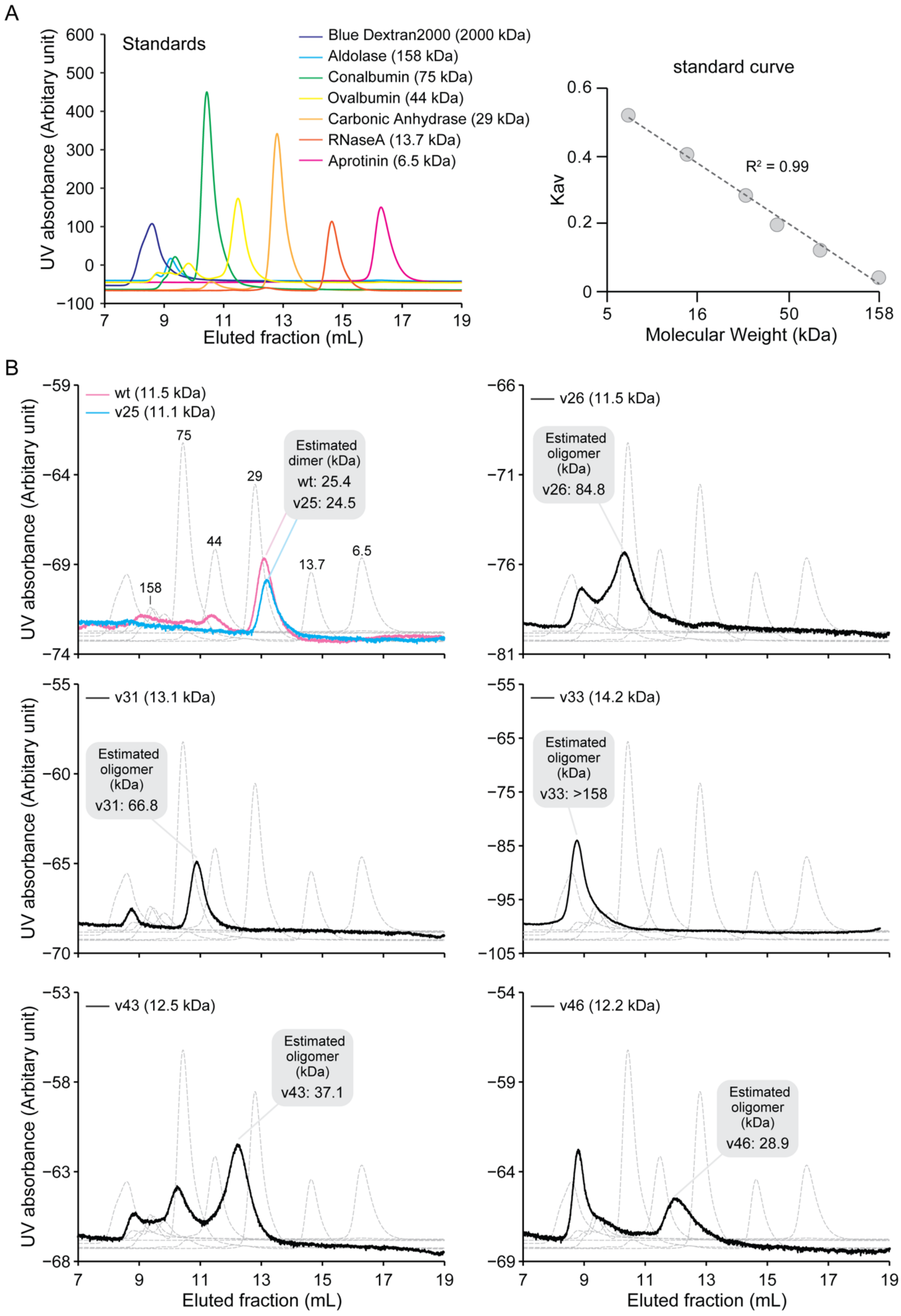

A
Standards
Blue Dextran2000 (2000 kDa)
Aldolase (158 kDa)
Conalbumin (75 kDa)
Ovalbumin (44 kDa)
Carbonic Anhydrase (29 kDa)
RNaseA (13.7 kDa)
Aprotinin (6.5 kDa)
UV absorbance (Arbitary unit)
Eluted fraction (mL)
standard curve
R² = 0.99
Kav
Molecular Weight (kDa)
B
wt (11.5 kDa)
v25 (11.1 kDa)
Estimated dimer (kDa)
wt: 25.4
v25: 24.5
v26 (11.5 kDa)
Estimated oligomer (kDa)
v26: 84.8
v31 (13.1 kDa)
Estimated oligomer (kDa)
v31: 66.8
v33 (14.2 kDa)
Estimated oligomer (kDa)
v33: >158
v43 (12.5 kDa)
Estimated oligomer (kDa)
v43: 37.1
v46 (12.2 kDa)
Estimated oligomer (kDa)
v46: 28.9

**Fig. S16. Size-exclusion chromatography of synMinE variants.** (**A**) Eluted fraction of size-exclusion chromatography for standard proteins and the standard curve. (**B**) Eluted fraction of synMinE variants. synMinEv25 indicates similar eluted peak to the wildtype MinE, showing its proper dimerization. On the other hand, other synMinE variants show distant eluted peaks from wildtype and several peaks within the same variants are also observed, suggesting that oligomers of those variants are bigger than the dimer or tend to be aggregated.

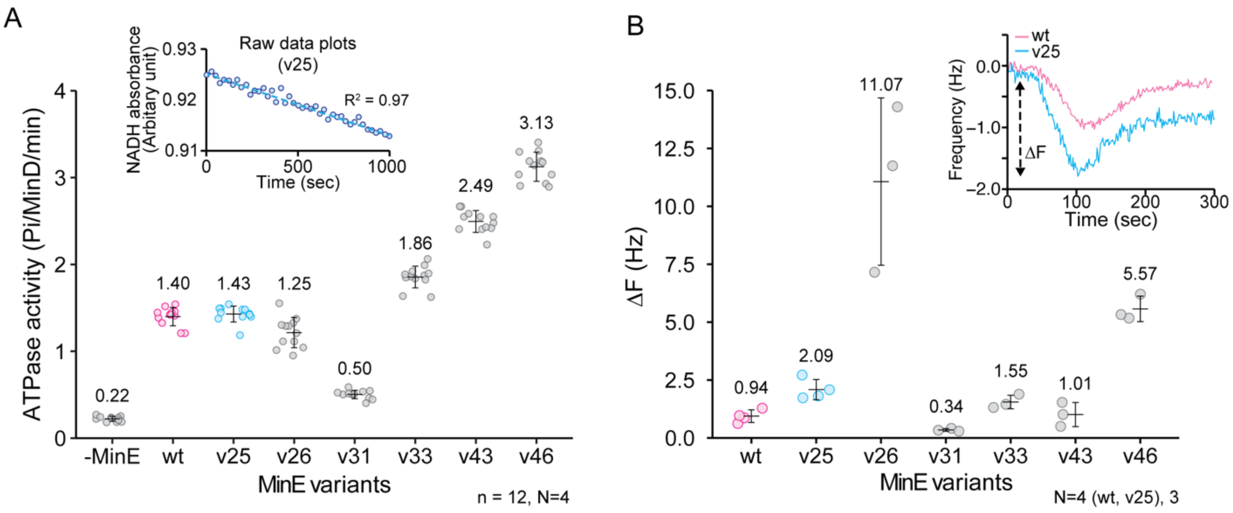


**Fig. S17. ATPase activity and QCMD assay.** (**A**) An ATPase assay reveals that all tested synMinE variants induce MinD's ATPase activity, although induced ATPase activities vary between 36% (v31) and 226% (v46) compared to wildtype, suggesting ATPase induction has to be finely tuned for proper function of MinDE oscillation. The subset shows the raw data plots of ATPase measurement with synMinEv25. (**B**) QCMD measurements indicate that all tested synMinE variants bind to the model lipid membrane, although binding strength is highly variying, depending on variants. However, Fig. 4D suggests that the difference in membrane binding may not be related to the cell phenotype. The subset shows the comparison of raw QCMD measurement between wt and v25.

**Table S1. DNA sequences and primers used in this study.**

| Name | Sequence (5' to 3') |
|---|---|
| 5' additional sequence for synMinE library | CCCGCGAAATTAATACGACTCACTATAGGG AGACCACAACGGTTTCCCTCTAGAAATAAT TTTGTTTAACTTTAAGAAGGAGATATACCAT G |
| 3' additional sequence for synMinE library | TAACTAGCATAACCCCTTGGGGCCTCTAAA CGGGTCTTGAGGGGTTTTTTG |
| MinD Ins3_FW (for linearization of pMLB-sfGFP-MinD.MinE) | GAATTCGCACGCATTATTGTTG |
| pMLB-lin-RV (for linearization of pMLB-sfGFP-MinD.MinE) | ATGTATATCTCCTTCTTAAATCTAGA |
| mGreenLantern-opt-FW (for insertion of mGreenLantern gene) | AGATATACATATGGTTAGTAAAGGAGAAGA AT |
| MinDuElin-RV (for insertion of mGreenLantern gene) | GAATTCTTTGTAGAGCTCATC |
| pMLB-ENDlin-FW (for linearization of the pMLB plasmid for synMinE genes) | GCCCGCTGTAAAAGCGCA |
| MinEdel2-12-RV (for linearization of the pMLB plasmid for synMinE genes) | CATAACTTATCCTCCGA |
| synMinEv5-OHminD-FW (for insertion of synMinEv5) | AGGATAAGTTATGAGTATTTTAGATTTTTTC TTTCCTTC |
| synMinEv5-OHpMBL-RV (for insertion of synMinEv5) | TACAGCGGGCTTATTGTTGTTCAGGTAATTG GATG |
| synMinEv10-OHminD-FW (for insertion of synMinEv10) | AGGATAAGTTATGTCATTTTTATCTTATTTA TTTGGTG |
| synMinEv10-OHpMBL-RV (for insertion of synMinEv10) | TACAGCGGGCTTAAGTAAGTTCCTGCTCAG ATAAC |
| synMinEv25-OH-FW (for insertion of synMinEv25) | AGGATAAGTTATGTCAATTTTTGATTTTTTT ACTGC |
| synMinEv25-OH-RV (for insertion of synMinEv25) | CAGCGGGCTTAGGCTTGCGCGGGAGCCTC |
| synMinEv26-OHminD-FW (for insertion of synMinEv26) | AGGATAAGTTATGTCAATTTTTGATTATTTT AAATC |
| synMinEv26-OHpMBL-RV (for insertion of synMinEv26) | TACAGCGGGCTTACAGAGACTGTTCTGGCA G |
| synMinEv29-OHminD-FW (for insertion of synMinEv29) | AGGATAAGTTATGATGATGGGTGAATTTAT TAGTCG |
| synMinEv29-OHpMBL-RV (for insertion of synMinEv29) | TACAGCGGGCTTACTGTTCGGCCGCGTAAT C |
| synMinEv31-OHminD-FW (for insertion of synMinEv31) | AGGATAAGTTATGATGTTAGCTGAATTTATT AATATG |

**Table S1. DNA sequences and primers used in this study (continued).**

| Name | Sequence (5' to 3') |
|---|---|
| synMinEv31-OHpMBL-RV (for insertion of synMinEv31) | TACAGCGGGCTTAAGATGTCTCGCTGGGCGA |
| synMinEv33-OHminD-FW (for insertion of synMinEv33) | AGGATAAGTTATGATGGGTGAATTTATTTCAAAAATG |
| synMinEv33-OHpMBL-RV (for insertion of synMinEv33) | TACAGCGGGCTTACTGGGGCACAGTTAACGC |
| synMinEv35-OHminD-FW (for insertion of synMinEv35) | AGGATAAGTTATGTTAATTATGTCAATTTTATCATTTTTTCG |
| synMinEv35-OHpMBL-RV (for insertion of synMinEv35) | TACAGCGGGCTTATGCCGGTGTTGTTGGCG |
| synMinEv37-OHminD-FW (for insertion of synMinEv37) | AGGATAAGTTATGAATATTTTACAATTGTTTAGTCGTAC |
| synMinEv37-OHpMBL-RV (for insertion of synMinEv37) | TACAGCGGGCTTAGGCTCCTAAATCCTCAAGTAC |
| synMinEv40-OHminD-FW (for insertion of synMinEv40) | AGGATAAGTTATGTCAATTTTAGGAATTTTTC |
| synMinEv40-OHpMBL-RV (for insertion of synMinEv40) | TACAGCGGGCTTATGCTGCTTTTGGTTTAGACGC |
| synMinEv43-OHminD-FW (for insertion of synMinEv43) | AGGATAAGTTATGTCAATTTTAGATTATTTCTTTTCAAG |
| synMinEv43-OHpMBL-RV (for insertion of synMinEv43) | TACAGCGGGCTTACGGCGGGGCTTGTGGTGG |
| synMinEv44-OHminD-FW (for insertion of synMinEv44) | AGGATAAGTTATGTCATTTTTAGCCTTTTTCTTTG |
| synMinEv44-OHpMBL-RV (for insertion of synMinEv44) | TACAGCGGGCTTAAGACTGCTCCGGCAGAGT |
| synMinEv46-OHminD-FW (for insertion of synMinEv46) | AGGATAAGTTATGATGTTTCCATTAATTGATAAATTAT |
| synMinEv46-OHpMBL-RV (for insertion of synMinEv46) | TACAGCGGGCTTAAGGTTCTGTCGACGGCGT |
| synMinEv48-OHminD-FW (for insertion of synMinEv48) | AGGATAAGTTATGTTTGGATTTAGTTTTCG |
| synMinEv48-OHpMBL-RV (for insertion of synMinEv48) | TACAGCGGGCTTAGGAACTCAAAGCGGAGGG |
| MinD-RV (to obtain pMLB-mGreenLantern-MinD, used with pMLB-ENDlin-FW primer) | TTATCCTCCGAACAAGCG |
| mGL-opt-Stop-RV (to obtain pMLB-mGreenLantern, used with pMLB-ENDlin-FW primer) | TATTTGTAGAGCTCATCCATGTCATGTG |

**Table S1. DNA sequences and primers used in this study (continued).**

| Name | Sequence (5' to 3') |
|---|---|
| Linker-His-FW (for linearization of pET28a plasmid) | GGTGGATCTGGAGTCGAGC |
| del_MinE_rev (for linearization of pET28a plasmid) | CATGGTATATCTCCTTCTTAAAGTTAA |
| synMinEv5-OHp28a-FW (for insertion of synMinEv5 in pET28a) | GATATACCATGAGTATTTTAGATTTTTTCTTTCCTTC |
| synMinEv5-OHp28a-RV (for insertion of synMinEv5 in pET28a) | TCCAGATCCACCTTGTTGTTCAGGTAATTGGATG |
| synMinEv10-OHp28a-FW (for insertion of synMinEv10 in pET28a) | GATATACCATGTCATTTTTATCTTATTTATTTGGTG |
| synMinEv10-OHp28a-RV (for insertion of synMinEv10 in pET28a) | TCCAGATCCACCAGTAAGTTCCTGCTCAGATAAC |
| synMinEv25-OHp28a-FW (for insertion of synMinEv25 in pET28a) | GATATACCATGTCAATTTTTGATTTTTTTACTGC |
| synMinEv25-OHp28a-RV (for insertion of synMinEv25 in pET28a) | TCCAGATCCACCGGCTTGCGCGGGAGCCTC |
| synMinEv26-OHp28a-FW (for insertion of synMinEv26 in pET28a) | GATATACCATGTCAATTTTTGATTATTTTAAATC |
| synMinEv26-OHp28a-RV (for insertion of synMinEv26 in pET28a) | TCCAGATCCACCCAGAGACTGTTCTGGCAG |
| synMinEv29-OHp28a-FW (for insertion of synMinEv29 in pET28a) | GATATACCATGATGATGGGTGAATTTATTAGTCG |
| synMinEv29-OHp28a-RV (for insertion of synMinEv29 in pET28a) | TCCAGATCCACCCTGTTCGGCCGCGTAATC |
| synMinEv31-OHp28a-FW (for insertion of synMinEv31 in pET28a) | GATATACCATGATGTTAGCTGAATTTATTAATATG |
| synMinEv31-OHp28a-RV (for insertion of synMinEv31 in pET28a) | TCCAGATCCACCAGATGTCTCGCTGGGCGA |
| synMinEv33-OHp28a-FW (for insertion of synMinEv33 in pET28a) | GATATACCATGATGGGTGAATTTATTTCAAAAATG |
| synMinEv33-OHp28a-RV (for insertion of synMinEv33 in pET28a) | TCCAGATCCACCCTGGGGCACAGTTAACGC |
| synMinEv43-OHp28a-FW (for insertion of synMinEv43 in pET28a) | GATATACCATGTCAATTTTAGATTATTTCTTTTCAAG |
| synMinEv43-OHp28a-RV (for insertion of synMinEv43 in pET28a) | TCCAGATCCACCCGGCGGGGCTTGTGGTGG |
| synMinEv44-OHp28a-FW (for insertion of synMinEv44 in pET28a) | GATATACCATGTCATTTTTAGCCTTTTTCTTTG |
| synMinEv44-OHp28a-RV (for insertion of synMinEv44 in pET28a) | TCCAGATCCACCAGACTGCTCCGGCAGAGT |

**Table S1. DNA sequences and primers used in this study (continued).**

| Name | Sequence (5' to 3') |
|---|---|
| synMinEv46-OHp28a-FW (for insertion of synMinEv46 in pET28a) | GATATACCATGATGTTTCCATTAATTGATAAATTAT |
| synMinEv46-OHp28a-RV (for insertion of synMinEv46 in pET28a) | TCCAGATCCACCAGGTTCTGTCGACGGCGT |

**Movie S1.**

Min oscillations induced by synMinEv25 inside lipid droplets as a representative "positive" variant found in the *in vitro* screening. Cell-free expressed synMinEv25 was encapsulated in lipid droplets with 1 μM EGFP-MinD, 2.5 mM ATP, and 10 g/L BSA. Then, synMinEv25 self-assembled into spatiotemporal patterns on the membrane together with MinD (shown in green). Timestamp indicates mm:ss. Scale bar: 20 μm.

**Movie S2.**

Homogeneous membrane binding of EGFP-MinD (shown in green) together with synMinEv4 inside lipid droplets as a representative "negative" variant found in the *in vitro* screening. Cell-free expressed synMinEv4 did not induce any dynamics behavior of Min proteins inside lipid droplets. Timestamp indicates mm:ss. Scale bar: 20 μm.

**Movie S3.**

Min oscillations inside Δ*min*DE *E. coli* cells transformed with synMinEv25 and mGreenLantern-MinD. synMinEv25 fulfills the emergent function of Min proteins for cell division and therefore gives normal cell phenotype and their oscillatory movement. Timestamp indicates mm:ss. Scale bar: 10 μm.

**Movie S4.**

Zoomed-in view of *E. coli* cells showing Min oscillations induced by synMinEv25 and mGreenLantern-MinD. Timestamp indicates mm:ss. Scale bar: 2 μm.

**Movie S5.**

Min oscillations inside minicell phenotype *E. coli* cells induced by synMinEv5. Minicells can be observed as "spots" together with rod-shaped cells containing Min oscillations (mGreenLantern-MinD is shown in green), indicating synMinEv5 induces Min oscillations but cannot lead to proper cell division. Timestamp indicates mm:ss. Scale bar: 2 μm.

**Movie S6.**

Min oscillations inside filamentous cells transformed with synMinEv37 and mGreenLantern-MinD (shown in green). Even inside the division-defect cells Min oscillations can emerge, hence, some synMinE variants can function as an inducer of Min protein dynamics, but not as a proper cell division machinery. Timestamp indicates mm:ss. Scale bar: 20 µm.

**Movie S7.**

Min oscillations inside Δ*min*DE *E. coli* cells transformed with wtMinE and mGreenLantern-MinD as a positive control. Timestamp indicates mm:ss. Scale bar: 10 µm.

**Data S1. (separate file)**

Amino acid and DNA sequences of all synMinE variants used in this study, together with similarity and identity scores to *E. coli* MinE and its closest homolog (with accession number of homologs). Also, *in silico* ranking, *in vitro* screening result, *in vivo* phenotype, length, and estimated molecular weight are included.
